# Machine learning reveals sequence and genomic context features underlying *Alu*-specific effects on genome folding

**DOI:** 10.64898/2026.09.16.752217

**Authors:** Shu Zhang, Katherine S. Pollard

## Abstract

The *Alu* transposable element is among the most abundant classes of mobile DNA in the human genome and has been linked to gene regulation and chromatin organization. Yet how individual *Alu* insertions influence nearby chromatin interactions remains poorly understood. To investigate this, we used deep learning to perform a genome-wide *in silico* deletion screen of ∼1.1 million *Alus*, predicting each element’s importance to local genome folding. We identified a subset of high-scoring *Alus* that span multiple *Alu* subfamilies and are enriched in loci that are fast-evolving and gene-dense, especially loci encoding genes that are actively transcribed and/or related to *Alu* biology. We further found that polymorphic *Alus* preferentially occur in regions tolerant of sequence variation but predicted to be resistant to changes in chromatin structure. Targeted *in silico* mutagenesis showed that the importance of individual *Alus* to local chromatin interactions depends on both intrinsic *Alu* sequence properties and genomic context. Finally, we identified *Alu* sequences predicted to alter CTCF-mediated boundary strength and, in some cases, to promote the formation of new loops and boundaries. Together, these results position *Alus* as key modulators of genome architecture, while underscoring that the fate of a new *Alu* depends on where it inserts and how it interacts with other determinants of chromatin state and structure.

## Introduction

Repetitive elements (REs) occupy more than half of the human genome and have played a central role in shaping genome evolution and regulation. Historically considered as ‘junk DNA’, these elements are now recognized as widespread contributors to gene regulation, influencing transcription, chromatin accessibility, and epigenetic modifications ^1^. More recently, their role has expanded to include the folding of the genome into three-dimensional (3D) structures. This genome organization supports essential cellular processes and brings genes into proximity with their regulatory elements, enabling proper cell function and development. While key proteins such as CTCF and cohesin drive loop extrusion and large-scale chromatin structure, these factors alone do not fully explain the chromatin interactions we observe across different cell types and species ^2–4^. Identifying additional sequence-encoded determinants of genome folding is a key step towards understanding how dysregulation in genome organization contributes to disease or evolutionary differences.

Transposable elements (TEs) have recently emerged as underappreciated contributors to three-dimensional genome organization, although the mechanisms through which they act remain incompletely understood ^5–9^. Among these, *Alus* are of particular interest. Unlike most TE families, *Alus* are primate-specific, actively contributing to genetic variation today, and make up around 11% of the human genome ^10,11^. A*lus* are categorized into subfamilies of varying evolutionary age and sequence divergence, which allows us to compare how different *Alu* types contribute to gene regulation. Experimentally, *Alus* have been shown to host binding sites for regulatory proteins, mediate enhancer-promoter interactions, and help form species-specific chromatin loops ^5,12–14^. Consistent with these observations, REs, especially certain *Alus*, were predicted to alter local chromatin interactions when deleted *in silico* ^15^. Some *Alu* subfamilies were enriched for disruptive elements, but even these subfamilies included many elements predicted to be inert with respect to chromatin interactions. Thus, it remains to be determined what features of *Alus* and/or the genomic loci in which they insert determine whether any given *Alu* contributes to chromatin organization.

Since many *Alus* predicted to affect 3D chromatin are devoid of CTCF binding motifs and transcriptionally active regions ^15^, it appears that *Alus* modulate genome folding outside of well-characterized mechanisms. Investigating this question is challenging, because it is experimentally infeasible to systematically identify which *Alus* and which sequence features of them are important for chromatin organization. We therefore take advantage of recent advances in deep learning, where models trained on chromosome conformation capture data enable accurate predictions of 3D genome organization directly from DNA sequence ^16–18^. These models, combined with high throughput *in silico* mutagenesis (ISM) experiments ^19,20^, provide a powerful framework to systematically dissect how specific DNA sequence elements may influence chromatin interactions at scale.

Here, we leveraged the Akita deep learning model ^21^ to test the hypothesis that specific sequence features of *Alus* influence genome folding. We performed a genome-wide in silico deletion screen of ∼1.1 million *Alus* in the human genome to identify elements with the strongest predicted importance to local chromatin interactions. We detected an interplay between genomic context and intrinsic sequence properties: high-scoring *Alus* were enriched in gene-dense, active regions and in rapidly evolving loci. Further ISM experiments suggested that *Alus* may influence CTCF-mediated boundary regulation, modulating existing boundaries and occasionally promoting new predicted chromatin interactions. To explore *Alu* evolution, we analyzed polymorphic *Alus,* revealing genomic regions tolerant of sequence variation but resistant to changes in chromatin organization as hotspots for recent *Alu* insertions. Collectively, these analyses support *Alus* as modulators of 3D genome architecture, providing a mechanism by which TE variation can contribute to regulatory diversity and genome evolution.

## Results

### Machine learning predicts genome-wide *Alu* disruption to 3D genome folding

To systematically quantify the contributions of individual *Alu* elements to genome folding, we deleted every *Alu* in the hg38 reference human genome (n=1,109,106 post-filtering) and predicted how local chromatin interactions changed using the SuPreMo-Akita framework ^20^ (**Figure 1a**, **Methods**). The resulting “disruption scores” measure how important the *Alu* is to chromatin organization within the surrounding megabase (Mb), using the mean squared error (MSE) and one minus the Spearman correlation (CORR), two complementary metrics ^22^. Larger values indicate greater contribution of the deleted sequence to predicted chromatin contacts.

**Figure 1.**
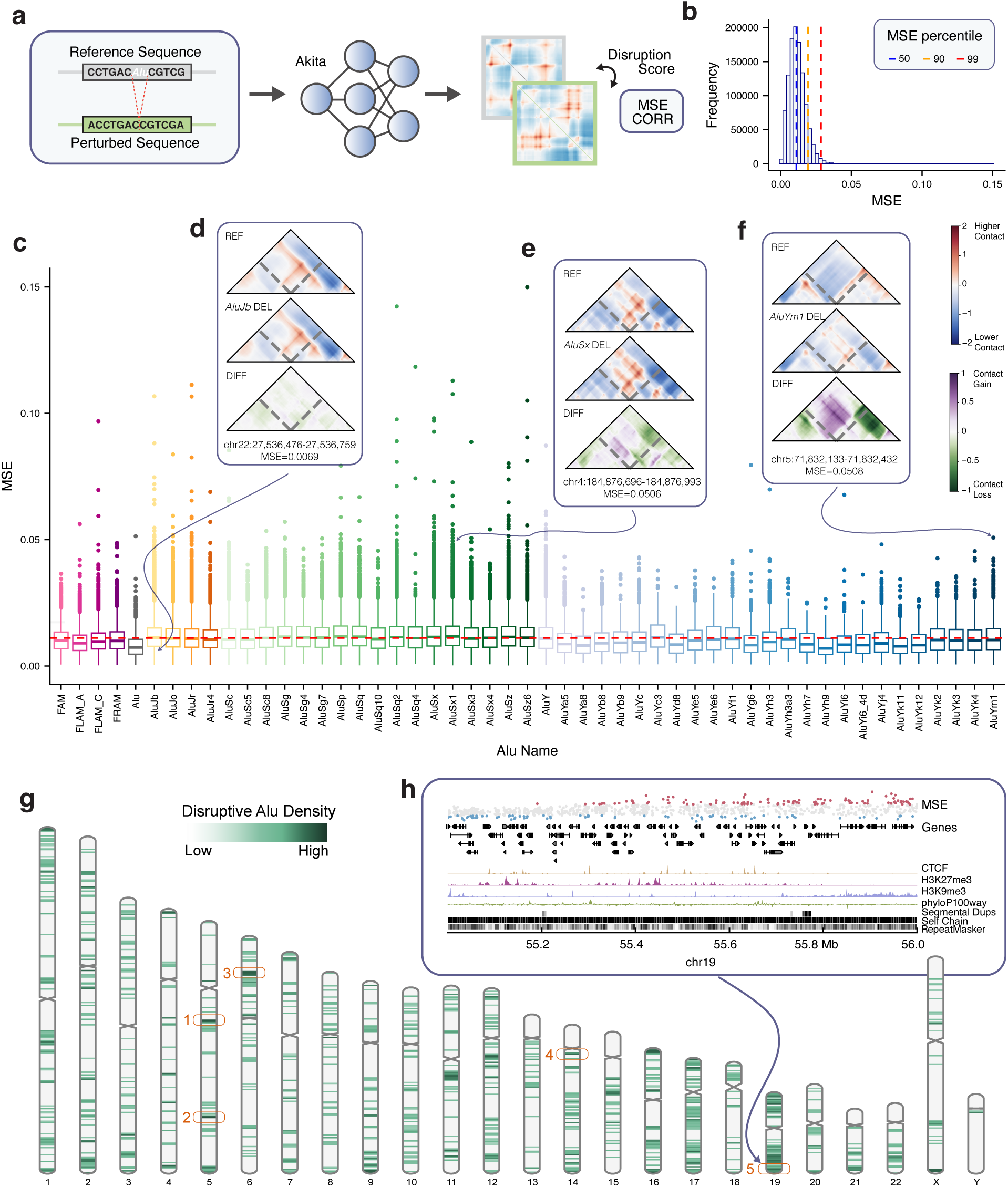
Predicted genome-wide *Alu*-associated chromatin folding. **a.** Scores for all hg38 *Alus* were calculated with SuPreMo-Akita by comparing Akita-predicted contact maps for unperturbed (REF) and *Alu*-deleted (ALT) sequences for the human foreskin fibroblast (HFF) cell type. MSE and CORR were used to quantify the change between the two maps (‘disruption’). **b.** Distribution of MSE scores, with dashed lines at the 50th, 90th, and 99th percentiles. *Alus* with MSE scores above the 99th percentile are defined as ‘highly disruptive’ when deleted, meaning Akita predicts that local chromatin interactions depend upon the *Alu*. **c.** Distribution of MSE scores across all *Alus*, grouped by the *Alu* type. Colors represent the broader *Alu* family each *Alu* belongs to. The dashed red line represents the median score across all *Alus*. **d–f.** Predicted contact maps for a neutral *AluJb* deletion (chr22, MSE=0.0069) and two highly disruptive deletions: *AluSx1* (chr4, MSE=0.0506), showing a strong change in contact intensity, and *AluYm1* (chr5, MSE=0.0508), showing altered chromatin structure. **g.** Density of highly disruptive *Alus* in 1 Mb bins genome-wide, normalized by total *Alu* count per bin. Numbered regions: (1) BDP1 (chr5: 71000000-72000000); (2) PDCH genes (chr5:141000000-14200000); (3) BTN and histone genes (chr6:25000000-28000000); (4) T-cell receptor alpha genes (chr14:21000001-2200000); (5) ZNF genes (chr19:58000001-58617616). **h.** Genome browser track (chr19:55,000,000–56,000,000), with individual *Alus* plotted as dots colored by score (blue = neutral, red = highly disruptive).

We averaged scores across six sequence augmentations to improve stability (**Methods**) and compared deletions with duplications and inversions as alternative perturbations. Deletions and duplications produced highly correlated disruption scores (Pearson *R* = 0.863), while inversions produced minimal effects, indicating that disruption is largely independent of *Alu* orientation and supporting deletions as a reproducible measure of *Alu* disruption (**Supplementary Figure 1a**).

Across all *Alu* deletions, the average disruption score was modest (mean MSE = 0.0117, median 0.0111, std 0.0058; **Figure 1b-c, Supplementary Table 1**), indicating that most individual *Alus* are not predicted to play a major role in local chromatin interactions, consistent with prior observations using a much smaller number of *Alus* ^15^. We found these trends to be consistent when scoring with CORR (mean CORR=0.0484, median 0.0370, std 0.0392, **Supplementary Figure 1b-d**). Within each family, individual outliers had markedly higher disruption than the typical *Alu*. Thus, we defined a ‘highly disruptive’ *Alu* as an *Alu* with an MSE disruption score at or above the 99th percentile (MSE≥0.0285), and compared them to ‘neutral’ *Alus*, or *Alus* with MSEs at or below the 50th percentile (MSE≤0.0111) (**Figure 1b, d-f**).

We observed slight differences in disruption score distributions across *Alu* families, with *AluSx1* and *AluYh9* showing the highest and lowest mean disruption, respectively (mean MSE=0.0123 and 0.0083). However, several other young *Alu* families, including the *AluYb9*, *AluYh3a3*, *AluYf1*, *AluYe6*, and *AluYc3* families, were enriched for highly disruptive elements (**Supplementary Figure 2 a-c**). Thus, although *Alu* family age is associated with importance to chromatin interactions, substantial heterogeneity within families suggests that family identity alone does not explain which *Alus* were most disruptive. This motivated us to examine both the genomic contexts and intrinsic sequence properties that may distinguish highly disruptive *Alus*.

### Highly disruptive *Alus* cluster in evolutionarily dynamic regions enriched for gene expansions and pseudogenes

To investigate whether *Alus* that contribute to chromatin interactions are concentrated in certain genomic contexts, we identified the 1Mb regions genome-wide with the highest density of highly disruptive *Alus* (**Figure 1g**). The top loci overlapped gene families that have undergone recent expansions, including the protocadherin, KRAB zinc finger protein (KZFP), histone, and BTN families (**Figure 1g-h, Supplementary Figure 3a**). Several of these gene families are primate-specific or have experienced high evolutionary turnover. KZFPs, in particular, are thought to have expanded in response to the proliferation of *Alus* and other TEs to repress and silence these elements ^23^. Meanwhile, protocadherins evolved through repeated tandem gene duplications in vertebrates, where combinatorial expression of protocadherin genes increases neuronal diversity ^24^.

These observations indicate that *Alus* may play a larger role in chromatin organization in genomic regions that have experienced high evolutionary turnover and lineage-specific duplications. This association may be partly mechanistic: *Alus* and other repetitive elements are known to mediate structural variation, and their presence may itself have contributed to these expansions ^25,26^. These loci are not necessarily hotspots for *Alus* overall but rather show a higher density of disruptive *Alus* specifically (**Supplementary Figure 3b**) suggesting that rapidly evolving loci preferentially tolerate — or are shaped by — *Alus* with predicted effects on chromatin organization. Consistent with this, genome-wide, high-disruption *Alus* showed greater overlap with segmental duplications and self-chain regions (Mann-Whitney U *p* < 2.2×10⁻¹⁶ for all comparisons, **Supplementary Figure 3c**).

The genomic window with the greatest density of highly disruptive *Alus* contains the BDP1 locus (**Supplementary Figure 3a-b**), which encodes a core subunit of TFIIIB, which in turn recruits RNA Polymerase III. This is notable because *Alu* elements are transcribed by RNA Polymerase III, and their internal promoter elements are recognized by TFIIIIC, which guides TFIIIB. Previous studies have demonstrated that binding of TFIIIC to *Alus* alters histone acetylation and chromatin architecture ^27^, and that TFIII binding sites have been under accelerated evolution ^28^. More broadly, these findings support a model in which *Alu* elements with functional roles in chromatin organization preferentially persist near genes that both control and are influenced by their presence, linking their genome-folding potential to these co-evolving regions.

### Polymorphic *Alus* preferentially occur into genomic contexts predicted to tolerate structural perturbation

The enrichment of high-scoring *Alus* in rapidly evolving loci suggests that genomic context influences which *Alu* insertions persist over time and become integrated into chromatin organization (**Supplementary Figure 3c-d)**. We therefore asked whether polymorphic *Alus,* which are still segregating in human populations, would show evidence of ongoing selection. We analyzed polymorphic structural variants (SVs) identified from 1,019 long-read genomes in the 1000 Genomes Project ^29^, considering all biallelic insertions and deletions. TEs accounted for 33.34% of insertions and 4.53% of deletions, of which 74.41% and 82.40% were *Alus*, respectively (**Figure 2a**). Unlike non-TE polymorphic SVs, which concentrated at telomeric and sub-telomeric regions, polymorphic *Alus* were distributed more evenly across chromosome bodies (**Figure 2b, Supplementary Figure 4a**), consistent with ongoing retrotransposition.

**Figure 2.**
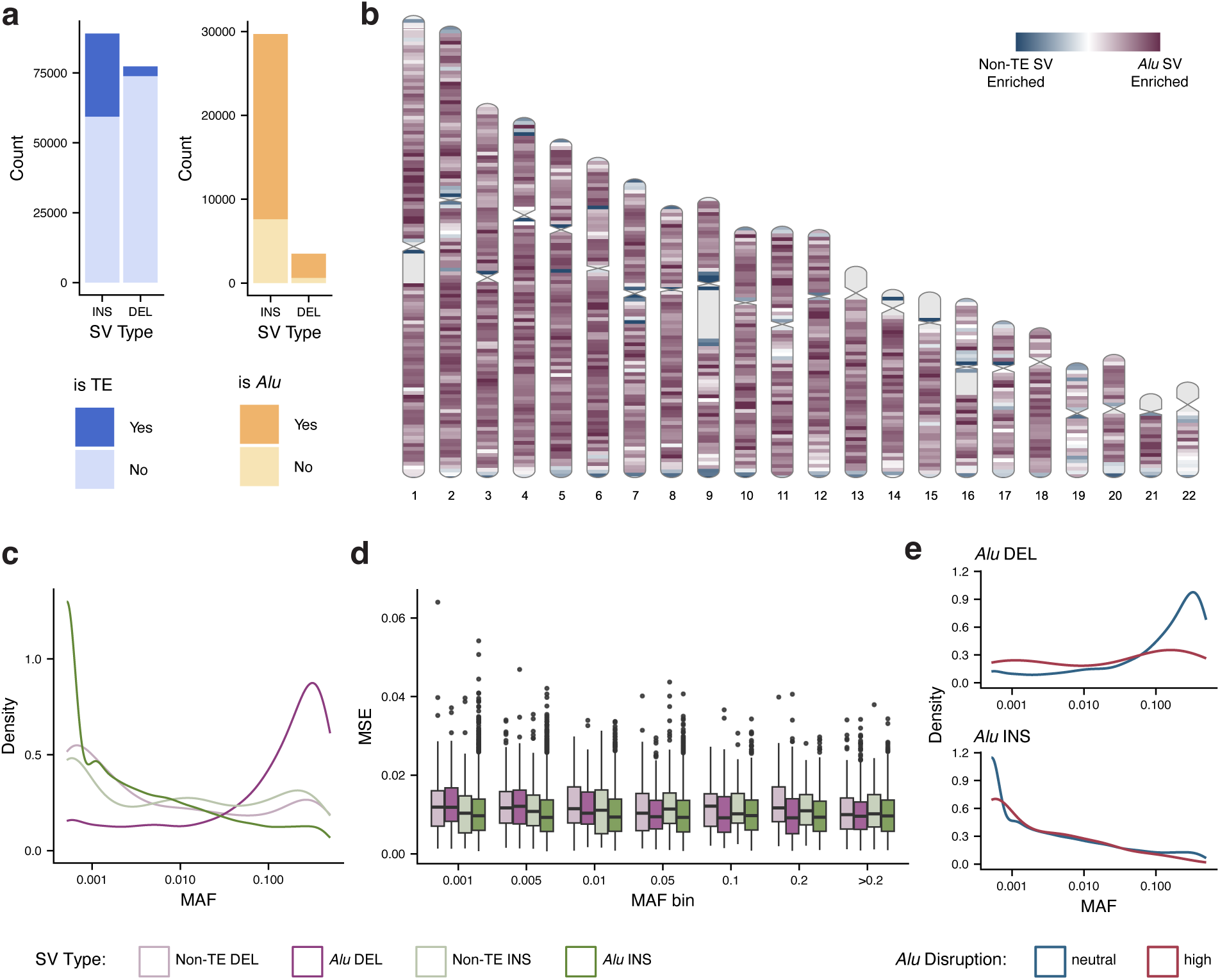
Predicted genome folding patterns of polymorphic *Alus* are consistent with purifying selection. **a.** Composition of biallelic, polymorphic SVs in the 1000 Genomes cohort. **b.** Karyoplot showing the enrichment of polymorphic SVs genome-wide, per 2Mb bin. Blue regions are non-TE SV-enriched, while mauve regions are *Alu*-enriched. **c.** MAF density (range 0-0.5) for polymorphic *Alu* insertions and deletions and length-matched non-TE SVs. **d.** Predicted disruption scores for *Alus* and length-matched non-TE SVs, by MAF bins. **e.** MAF density (range 0-0.5) for highly disruptive and neutral polymorphic *Alus,* across insertions and deletions.

Polymorphic *Alus* were modestly enriched in regions of high overall *Alu* density (Spearman ρ = 0.27 for insertions, ρ = 0.26 for deletions), but not in regions characterized by high-disruption *Alus* (**Supplementary Figure 4b-c**). Insertions and deletion frequencies in the same regions were moderately correlated, with a stronger correlation for non-TE SVs than *Alus* (**Supplementary Figure 4d**, Spearman ρ = 0.47 and 0.33, respectively).

We next examined allele frequencies of polymorphic *Alus*. At low minor allele frequencies (MAFs), *Alu* insertions were more common than non-TE insertions, while *Alu* deletions were enriched at higher MAFs (**Figure 2c**). This difference may partly reflect mutation processes, as ongoing *Alu* retrotransposition generates new insertions, which can contribute to their lower MAFs independently of selection. To ask whether selection on genome folding could contribute to these patterns, we scored all polymorphic SVs with SuPreMo-Akita and compared predicted disruption scores of polymorphic *Alu* insertions and deletions against those of length-matched non-TE SVs (**Supplementary Table 2, Methods**). Across both *Alu* and non-TE SVs, deletions were on average more disruptive than insertions, consistent with the greater potential for sequence loss to alter established chromatin interactions (**Figure 2d**). We then defined high-disruption (≥99th percentile MSE) and neutral (≤50th percentile MSE) thresholds from an independent genome-wide CHM13 *Alu* reference set (n = 1,060,253 post-filtering). High-disruption polymorphic *Alu* deletions had significantly lower MAFs than neutral deletions (mean MAF 0.107 vs. 0.197; Mann-Whitney U *p* = 0.008, Figure 2e). Although the corresponding comparison for insertions was not significant, disruptive *Alu* insertions still had a lower mean MAF than neutral insertions (mean MAF 0.015 vs. 0.030, *p* = 0.14). Together, these patterns suggest that the genomic consequences of *Alu* loss may influence the frequencies at which polymorphic deletions persist.

The relationship between polymorphism and predicted disruption also varied with *Alu* age. Polymorphic deletions were enriched for young *Alus*, specifically *AluYb9*, *AluYa5*, and *AluYb8* (**Supplementary Figure 5a, d-e,** ^30^). *In silico* deletions of these young polymorphic *Alus* were on average predicted to be less disruptive than deletions of fixed *Alus* within the same family (**Supplementary Figure 5b-c, Supplementary Table 3,** stratified Mann-Whitney U *p* = 2.6×10⁻^4^). By contrast, polymorphic deletions of older *Alus* were rarer but predicted to be more disruptive than fixed *Alu* deletions (*p* = 1.95×10⁻⁹). This pattern is consistent with two complementary processes — young *Alus* have had less time to become incorporated into functional chromatin environments, making their removal more tolerable, while older *Alus*, which tend to be more GC-rich and more frequently co-opted for functional purposes, may have greater consequences when deleted ^31,32, 33^.

### An *Alu’s* contribution to 3D chromatin is influenced by its genomic context

Given the enrichment of disruptive *Alus* in dynamic, expressed loci, we suspected that the surrounding genomic context of an *Alu* may help predict its potential to impact genome organization. We therefore trained logistic regression classifiers to separate highly disruptive and neutral *Alus* using genomic annotations across windows centered on each element, ranging from the *Alu* itself to the 1 Mb Akita prediction window (**Figure 3a, Supplementary Table 4**, **Methods**). Because many of these annotations are strongly correlated (**Supplementary Figure 6a**), we additionally fit models including pairwise interaction terms to capture combinatorial effects. Features calculated from the full 1Mb sequence achieved the strongest predictive

**Figure 3.**
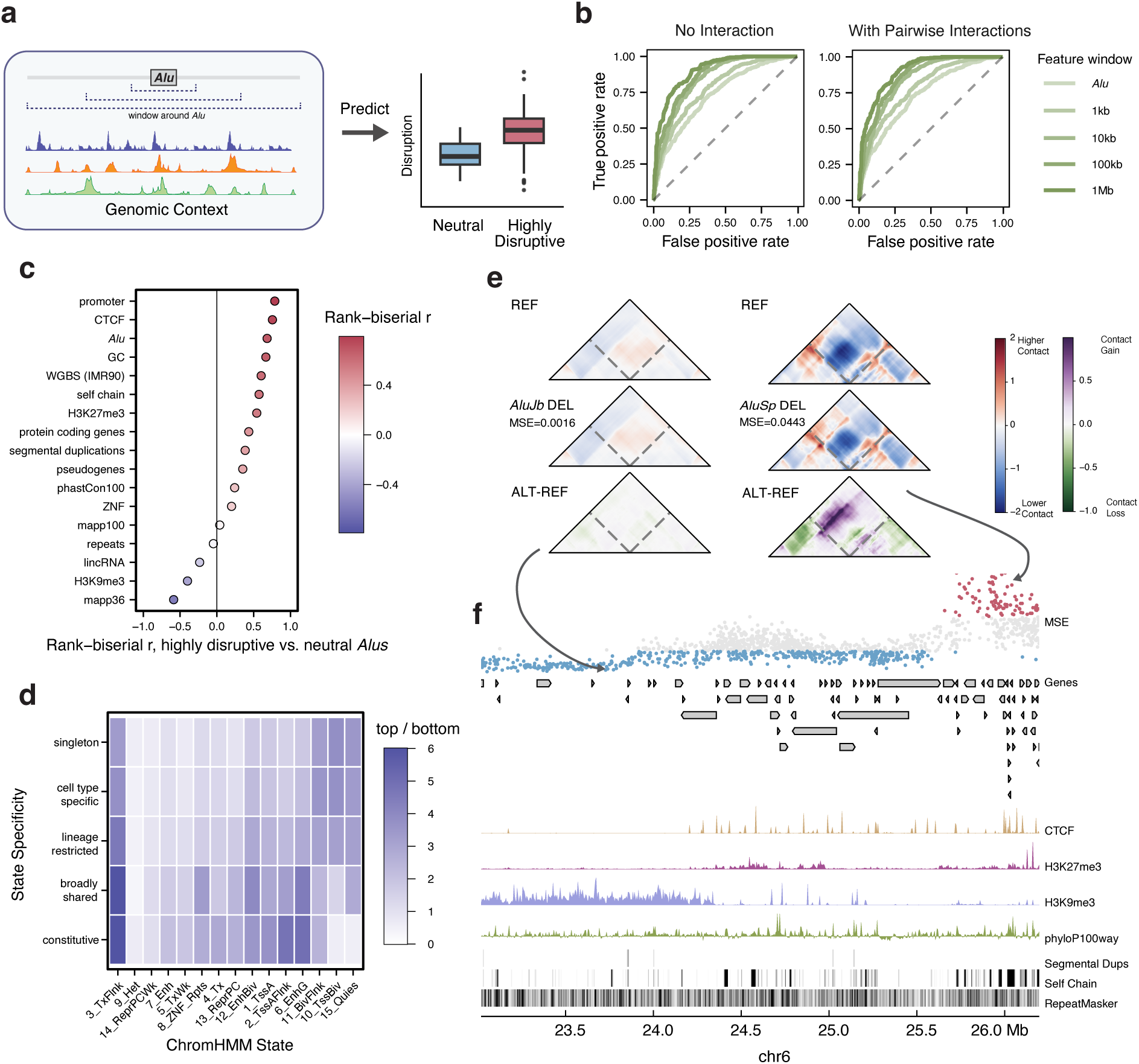
Genomic features predict *Alu*-associated genome folding. **a.** Schematic of the framework for predicting *Alu* disruption scores from genomic features. **b.** AUROC of logistic regression models classifying highly disruptive from neutral *Alus,* using features drawn from windows of increasing size around the *Alu* (*Alu* only, 1 kb, 10 kb, 100 kb, and 1 Mb). **c.** Rank-biserial *r* values for individual features at the 1Mb window. **d.** Enrichment of disruptive *Alus* across chromHMM states, grouped by the cell-type-specificity of each state. **e.** Predicted contact maps upon deletion of an *AluJb and AluSp,* drawn from low- and high-disruption genomic contexts, respectively, within the region below. **f.** Genome browser track (chr6:23000000-27000000) spanning a gene-sparse, heterochromatic region with low predicted *Alu* disruption next to a gene-rich, CTCF-bound region with high predicted disruption. Individual *Alus* are colored by predicted disruption level: blue (neutral), grey (intermediate), or red (highly disruptive).

power, reaching AUCs of 0.903 and 0.902 for the no-interaction and interaction models, respectively, on the held-out test set (**Figure 3b**). This substantially outperformed models restricted to local annotations at the *Alu* itself or smaller flanking regions, suggesting that the disruptive potential of an *Alu* is primarily determined by its broader genomic context. These results were consistent regardless of regression approach (**Supplementary Figure 7a**).

Among individual features, known determinants of genome folding were most associated with loci containing disruptive *Alus*, such as the number of CTCF binding sites, promoter density, DNA methylation levels and nearby *Alu* density (**Figure 3c, Supplementary Figure 7b, Methods**). All genomic features were derived from HFF cells, except for DNA methylation, for which IMR90 cells were used. Methylation and GC content became relatively more important at smaller windows (**Supplementary Figure 7b-c**), and methylation marks were more predictive when using genomic data from an embryonic stem cell line (H1ESC annotations, Akita predictions in HFF; **Supplementary Figure 7d-e**). We hypothesize that the difference in H1ESCs may reflect the known role of DNA methylation in silencing *Alus* and other TEs during early development ^34^. Overall, we find that broad genomic context is highly predictive of an individual *Alu’s* disruptive potential, while the relative contribution of individual features varies with genomic scale and cellular context.

To further characterize the genomic context associated with *Alu* disruption, we used ChromHMM annotations to compare chromatin environments enriched for highly disruptive *Alus* ^35^. We grouped each ChromHMM state into five categories based on cell-type specificity, ranging from constitutive to highly cell-type-specific regions. For each *Alu*, *we* calculated the proportion of its 1 Mb prediction window overlapping each category (**Methods**). Consistent with the regression results, disruptive *Alus* were enriched at actively transcribed regions and depleted in heterochromatin. The strongest enrichment was at constitutive State 3 regions, which mark transcription at the 5′ and 3′ ends of genes, and had 8.8-fold enrichment in highly disruptive *Alus* over neutral *Alus* (**Figure 3d**). This was followed by constitutive genic enhancers (State 6) and constitutive flanking regions of active transcription start sites (TSSs) (State 2), with 5-fold and 4.8-fold enrichment, respectively. Disruptive *Alus* were depleted at heterochromatin (State 9) across all categories, consistent with prior observations ^15^.

In some cases, we observed an inverse relationship between *Alu* density and cell-type-specificity of the ChromHMM state. In bivalent TSSs (State 10) and quiescent regions (State 15), disruptive *Alus* were depleted at constitutive states but became increasingly enriched with greater cell-type-specificity. For example, constitutive bivalent regions at evolutionarily conserved developmental regulators, such as *KLF14* and *FOXD3,* lacked disruptive *Alus* (**Supplementary Figure 8a-b**), whereas cell-type specific bivalent TSS regions at *AHNAK* and *CCDC155* harbored disruptive *Alus* (**Supplementary Figure 8c-e**). Many of these cell-type-specific bivalent regions were annotated in pluripotent lines, including the bivalent TSS of *CCDC155*, a meiosis-specific gene required for telomere attachment during gametogenesis.

Because retrotransposon integration occurs predominantly in the germline ^36^, *Alu* insertions near germline-restricted regulatory elements may be more likely to escape strong purifying selection and persist over evolutionary time. Their persistence could, in turn, provide opportunities for *Alus* to influence genome organization and regulatory activity at these loci ^37^.

### Disruptive *Alus* tend to be located in broadly expressed genes

The enrichment of disruptive *Alus* in gene-dense regions prompted us to examine genic *Alus* in greater detail. Of all scored *Alus*, 61% occurred within a gene; this proportion increased to 69% among highly disruptive *Alus*, consistent with our earlier findings that *Alus* in gene-dense regions tend to be more disruptive. The majority of these genes were protein-coding, with the remaining genes dominated by lncRNAs, lincRNAs, and pseudogenes (**Figure 4a**). Mean disruption among genic *Alus* was 0.0123, with substantial variation with gene categories.

**Figure 4.**
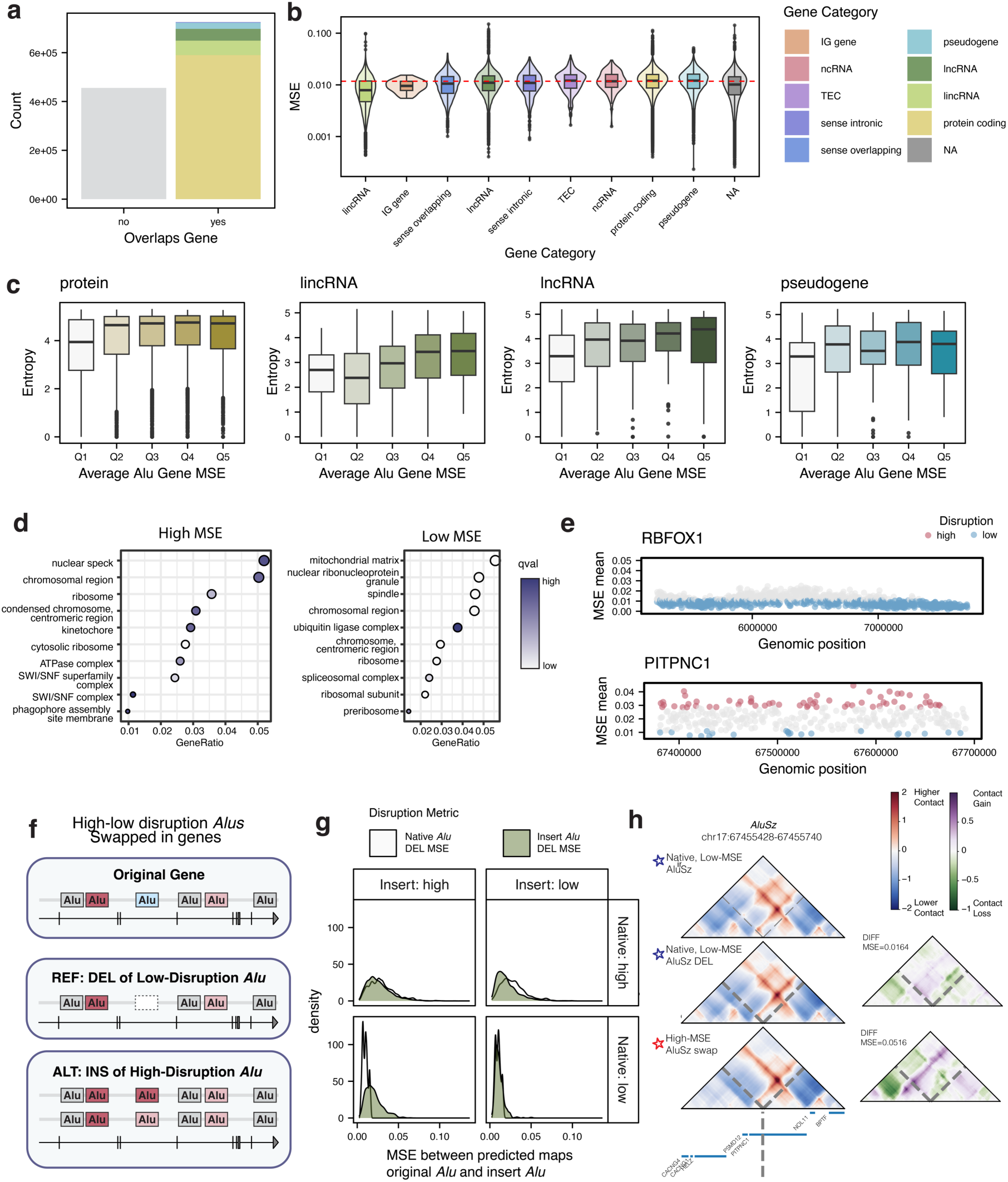
*Alus* predicted to influence chromatin interactions tend to be in broadly expressed genes. **a.** Proportion of scored *Alus* that overlap and do not overlap genes **b.** Distribution of MSE disruption scores for *Alus* located within genes. Colors represent the gene type in which the *Alu* is located. **c.** Average MSE of *Alu* deletions in a gene as a function of the expression entropy of the gene, averaged within the top four most prevalent gene categories. A higher entropy score indicates that the gene is expressed more broadly across cell types. **d.** Gene ontology comparisons for high-entropy genes with high average *Alu* scores (left, MSE > 80th percentile of genes) and low average *Alu* scores (right, MSE < 20th percentile of genes). **e.** Distributions of *Alu* scores within *RBFOX1*, a gene with high *Alu*-density but broadly low disruption, and *PITPNC1*, a gene containing many disruptive *Alus*, along with a wide range of disruption across the gene body. **f.** Framework for swapping high- and low-disruption *Alus* within a single gene. **g.** Predicted disruption across all four native/insert combinations of the *Alu*-swap experiment, aggregated over the top 25 genes by disruptive-*Alu* count. Within each combination, the white shading reflects disruption from the native *Alu’s* deletion, while the green shading reflects the disruption introduced by the swapped-in *Alu*, relative to that native-deletion baseline. **h.** Example locus in PITPNC1, where a low-MSE *AluSz* is swapped with a different low-MSE *AluSz*, along with a high-MSE *AluSz*.

We next asked whether genes harboring highly disruptive *Alus* showed differences in expression, using bulk RNA-seq from cell lines, primary cells, and in vitro differentiated cells.(**Methods, Supplementary Figure 9 a-c**). Across protein-coding genes, lncRNAs, lincRNAs, and pseudogene, *Alu* disruption was positively associated with expression across cell types, suggesting that broadly expressed genes tended to harbor more disruptive *Alus* (**Figure 4c**). This is consistent with our earlier observation that disruptive *Alus* were enriched at constitutive ChromHMM states associated with active transcription. Additionally, *Alu* disruption increased with the amount of functional constraint on the gene, even when controlling for the overall expression of that gene (**Supplementary Figure 9d**) ^38^.

Among broadly expressed genes, those with low *Alu* disruption were enriched for terms associated with core cellular processes, including translation, splicing, RNA processing, cell division, and mitochondrial function, using protein-coding genes as the background (**Figure 4d, Methods**). In contrast, genes with high *Alu* disruption were enriched for cellular machinery and functions linked to genome organization, most notably nuclear speckle localization and SWI/SNF chromatin remodeling (**Figure 4d**). This enrichment is consistent with a functional relationship between disruptive *Alus* and nuclear organization: recent evidence found that *Alu* RNAs organize actively transcribed genes surrounding nuclear speckles through interactions with genomic *Alus* ^39^. Our findings highlight a potential connection between disruptive *Alus* and genome organization at broadly expressed genes.

We next examined genes containing large numbers of *Alus*. As exemplified by *RBFOX1* and *CALN1*, many such genes tended to be lowly expressed, cell-type-specific, and embedded in heterochromatic regions. The *Alus* within these genes also had uniformly low disruption scores (**Figure 4e, Supplementary Figure 10a**). In contrast, genes like *PITPNC1* and *NGEF* contained *Alus* spanning a wide range of disruption scores (**Figure 4e**, **Supplementary Figure 10a)**, suggesting that individual *Alus* can have heterogeneous effects on local architecture within the same gene body. We found examples of this in both broadly expressed and more cell-type specific genes. Moreover, this co-occurrence of high-and low-scoring *Alus* within the same gene presented a way to disentangle the contributions of intrinsic *Alu* sequence and local genomic context.

We therefore performed *Alu*-swapping experiments across the top 25 genes with the largest number of high-scoring *Alus* (**Figure 4f, Supplementary Table 5, Methods**). For each pair of *Alus* within a gene, we exchanged sequences between neutral and highly disruptive *Alus* while keeping the surrounding genomic context unchanged; within-category swaps served as controls. Replacing a neutral *Alu* with a highly disruptive sequence substantially increased disruption at that locus (**Figure 4g**, rank-biserial effect size=0.699, Mann-Whitney U *p* < 2.2e-16), whereas the reciprocal swap produced a more modest decrease (effect size=-0.317, *p* < 2.2e-16). Thus, while *Alu* sequence contributes to disruption, local genomic context can sustain disruption even when the inserted sequence is less disruptive.

This sequence-dependent effect was evident in individual genes, including *PITPNC1*, where swapping a low-disruption *AluSz* with either another low-disruption or a high-disruption *AluSz* from the same gene produced correspondingly different disruption scores (**Figure 4h, Supplementary Figure 10b**). Insertion of the high-disruption *AluSz* was predicted to cause the loss of a loop-associated contact. Similar distributions were observed for *NGEF*, with representative contact maps illustrating the corresponding changes in local genome architecture (**Supplementary Figure 10b-c**). Across swaps, regions combining a high-disruption sequence with a high-disruption context maintained high disruption, whereas regions combining a neutral sequence with a neutral context remained minimally disruptive (**Figure 4g**). These findings indicate that the importance of individual *Alus* to chromatin organization is jointly shaped by intrinsic *Alu* sequence and its surrounding genomic context.

### Intrinsic *Alu* sequence features contribute to their disruption

Having shown that both *Alu* sequence and genomic context shape genome folding within variably sensitive genes, we next asked whether intrinsic sequence features contribute more broadly to *Alu* disruption scores genome wide. To test this, we compared the predicted effects of deleting *Alu* elements to deleting random, length-matched sequences *in silico*, retaining 90% or 99% of the surrounding genomic context (**Figure 5a, Supplementary Table 6, Methods**).

**Figure 5.**
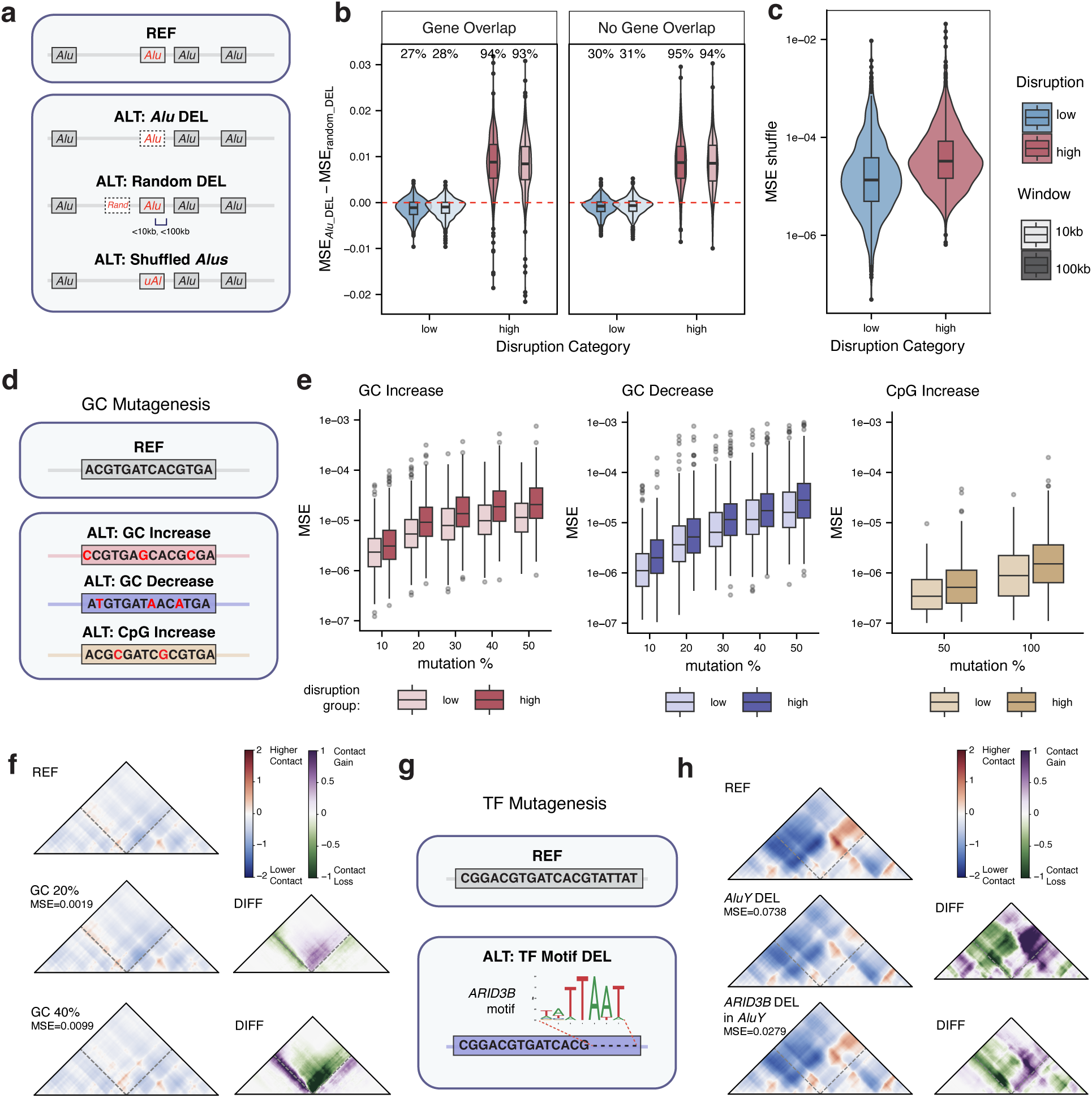
ISM of sequence-level features of *Alus* shape predicted disruption. **a.** Schematic of the ISM approach used to compare disruptive *Alus* to (1) randomly deleted, length-matched sequences and (2) shuffled *Alus.* **b.** Difference in prediction disruption for random deletions vs original *Alu* deletions, for *Alus* within and outside genes. The percentage of regions in which Alu deletion produced greater disruption than the corresponding neutral deletion is indicated. **c.** Predicted disruption scores for shuffled disruptive and neutral *Alus*. **d.** Schematic of *Alu* mutagenesis to alter GC and CpG content**. e.** Predicted disruption following increases in GC content, decreases in GC content, and increases in CpG content, respectively. **f.** Predicted contact maps surrounding a *FRAM* element (native GC=0.349) on chr19, following increases in GC content of 20% and 40%. **g.** Schematic of ISM of transcription factor motifs in *Alus*. **h.** Predicted contact maps following deletion of an *AluY* element or deletion of an *ARID3B* motif within the same *AluY* element on chr19.

Random deletions were further constrained to control for confounders: they avoided overlapping other *Alu* elements or annotated CTCF binding sites, and when the original *Alu* occurred within a gene, the matched deletion was restricted to genic regions.

For neutral *Alus*, deleting the *Alu* perturbed chromatin interactions slightly less than deleting a matched random sequence, consistent with Akita’s predictions in these cases being largely determined by the surrounding environment (Δ = −0.0011 MSE, paired Wilcoxon signed-rank *p* < 2.2×10⁻¹⁶, **Figure 5b**). In contrast, for highly disruptive *Alus*, deleting the *Alu* produced significantly greater disruption than deleting a random sequence (Δ = +0.0086 MSE, *p* < 2.2×10⁻¹⁶, **Figure 5b**), indicating that intrinsic sequence features contribute beyond genomic context. The difference between *Alu* and random-sequence perturbations was significantly greater for highly disruptive than neutral *Alus* (rank-biserial = 0.89, Mann-Whitney U *p* < 2.2×10⁻¹⁶), confirming that the intrinsic-sequence contribution is specific to disruptive *Alus*. We saw a similar pattern when shuffling *Alu* nucleotides (i.e., preserving base composition while changing subsequences): shuffling a highly disruptive *Alu* increased predicted disruption by 103% relative to shuffling a neutral *Alu* (**Figure 5c**, Mann-Whitney-U *p* < 2.2×10⁻¹⁶). This confirms that our findings are robust to the type of in silico perturbation used to assess sequence importance and supporting the idea that sequence features within disruptive *Alus* contribute to their effects.

### GC content and sequence motifs contribute to an *Alu’s* importance to 3D chromatin

If sequence composition of an *Alu* is associated with disruption, we hypothesized that targeted perturbations of these sequence features would produce measurable changes in predicted genome folding. Among the most informative features in our regression model were annotations associated with methylation and GC content, both of which can be readily perturbed *in silico* (**Figure 3c**, **Supplementary Figure 7a**). We therefore systematically increased or decreased the GC content in *Alus* and their flanking sequences. To simulate perturbing DNA methylation, we selectively introduced CpG dinucleotides while preserving overall sequence length and minimizing additional sequence changes (**Figure 5d, Supplementary Table 6, Methods**).

Both increasing and decreasing GC content produced measurable shifts in Akita’s predictions: across all mutation levels, increasing GC within the *Alu* body altered predicted chromatin interactions more than increasing GC in the flanks (Wilcoxon signed-rank *p* = 1.4×10⁻⁶– 1.0×10⁻²). Conversely, decreasing GC in the flanks produced larger shifts than in the body (Wilcoxon signed-rank *p* = 4.6×10⁻⁹–5.3×10⁻²; **Supplementary Figure 11a-b**). The predicted effect of changing GC content was also around 1.68 times higher for highly disruptive *Alus* across all levels and direction of GC mutation tested (**Figure 5e**, median fold-change 1.3×–2.0×, Mann-Whitney U *p* < 1.1×10⁻⁵ at every level), consistent with GC composition acting as a primary determinant of an *Alu’s* impact on folding rather than a passive correlate. Among *Alus* that were the most sensitive to altering GC content, we identified a case in a *FRAM* element with varying responses to GC alteration: a 20% increase in GC predicts strengthening an existing boundary, while a 40% increase weakens the same boundary but predicts formation of a new loop (**Figure 5f**). This suggests a dose-dependent relationship between GC content and boundary behavior, where moderate GC enrichment affects existing chromatin structures, while more extreme enrichment redirects contact formation. Separately, we hypothesize that lowering GC content raises the frequency of AT-rich motifs that lead to predicted disruption. For instance, nucleosomes are often displaced or excluded at AT-rich sequences^40,41^. We additionally tested CpG-content increases and found a similar pattern: high-disruption *Alus* showed a significantly larger response to CpG enrichment than low-disruption *Alus* (Mann-Whitney U *p* < 10⁻⁴⁶).

Because GC content is a coarse summary of sequence, we next asked whether specific binding motifs within *Alus* contribute to disruption beyond bulk composition. We found both *de novo* and known transcription factor binding motifs enriched in highly disruptive *Alus*, controlling for *Alu* family (**Supplementary Table 7, Methods**). Many of the known TF motifs were GC rich, consistent with the tendency of *Alus* to be in GC-rich regions. Enrichment of motifs from JASPAR and HOCOMOCO were assessed, and we found that zinc finger motifs were enriched across both databases. These include general GC-box binders such as the SP and KLF transcription factor families, as well as KZFPs such as ZNF263, which we know to engage with *Alus* and other TEs ^23,42^. In *AluYs*, the youngest subfamily, only the *ARID3B* motif was enriched, a transcription factor critical for embryonic development and potentially involved in regulating chromatin structure ^43^.

To test whether this enrichment reflected a causal relationship, we deleted the *ARID3B* motif from disruptive *AluY* elements. In most *AluYs,* motif deletion produced only modest changes in predicted disruption. However, in a few cases, including *AluY* elements on chr19 and chr9, deleting the motif predicted chromatin changes at similar loci affected by deletion of the entire *AluY*, although the effects were less pronounced in magnitude (**Figure 5h, Supplementary Figure 12a**). Thus, while the *ARID3B* motif can contribute to the predicted effect of an *AluY*, its presence alone does not generally explain *Alu* disruption. Consistent with this, gradients of the input sequences showed that the most relevant nucleotides were not localized to the *ARID3B* motif but were distributed broadly across the *Alu* body (**Supplementary Figure b-c**), suggesting that the contribution of the motif may depend on its surrounding sequence context.

We next asked whether other discrete sequence features within *Alus* showed similar sensitivity. No specific sub-sequence within the *Alu* sequence was predicted to be disproportionately sensitive to deletion within any of the *Alu* families (**Supplementary Figure 13a, Methods**).

Together, these results suggest that disruption generally reflects overall sequence content rather than individual TF binding sites or other discrete sequence features, with the notable exception of CTCF motifs. This is consistent with previous analyses and suggests that sequence features contributing to *Alu* sensitivity are not necessarily positionally constrained within the element^15,21^.

### *Alus* with roles in 3D chromatin contain intrinsic sequence features that alter predicted CTCF sensitivity

To better understand how sequence variation within *Alus* contributes differentially to genome folding, we asked whether different *Alus* produced different effects in the same genomic context. We used SuPreMo-Akita to predict chromatin interaction maps after substituting consensus sequences ^44^ from multiple *Alu* families in place of highly disruptive native *Alus* (**Figure 6a, Supplementary Table 6, Methods**), generating three versions of each locus: the native *Alu*, the deleted *Alu*, and the inserted consensus *Alu*. Across all loci, sequence similarity between the native *Alu* and substituted *Alu* was a poor predictor of contact map similarity; percent alignment was not correlated with disruption scores (Spearman ρ = 0.031, **Figure 6b**). Nevertheless, individual loci showed substantial variation in response to different *Alu* sequences. At an *AluSp* on chr5 (**Figure 6c**), related subfamilies tended to produce similar folding phenotypes, although sequence relatedness did not fully account for the observed variation. For example, the closely related *AluSc5* and *AluSg7* both produced strong disruption and predicted boundary formation, whereas the more distant *AluYk12* produced a contact map like that of the native *AluSp* (**Figure 6d-e**). A second example showing similar sequence-dependent variation is presented in **Supplementary Figure 14.** These results further support the conclusion that specific sequence features, beyond shared sequences within *Alu* families, determine how individual *Alus* influence genome folding.

**Figure 6.**
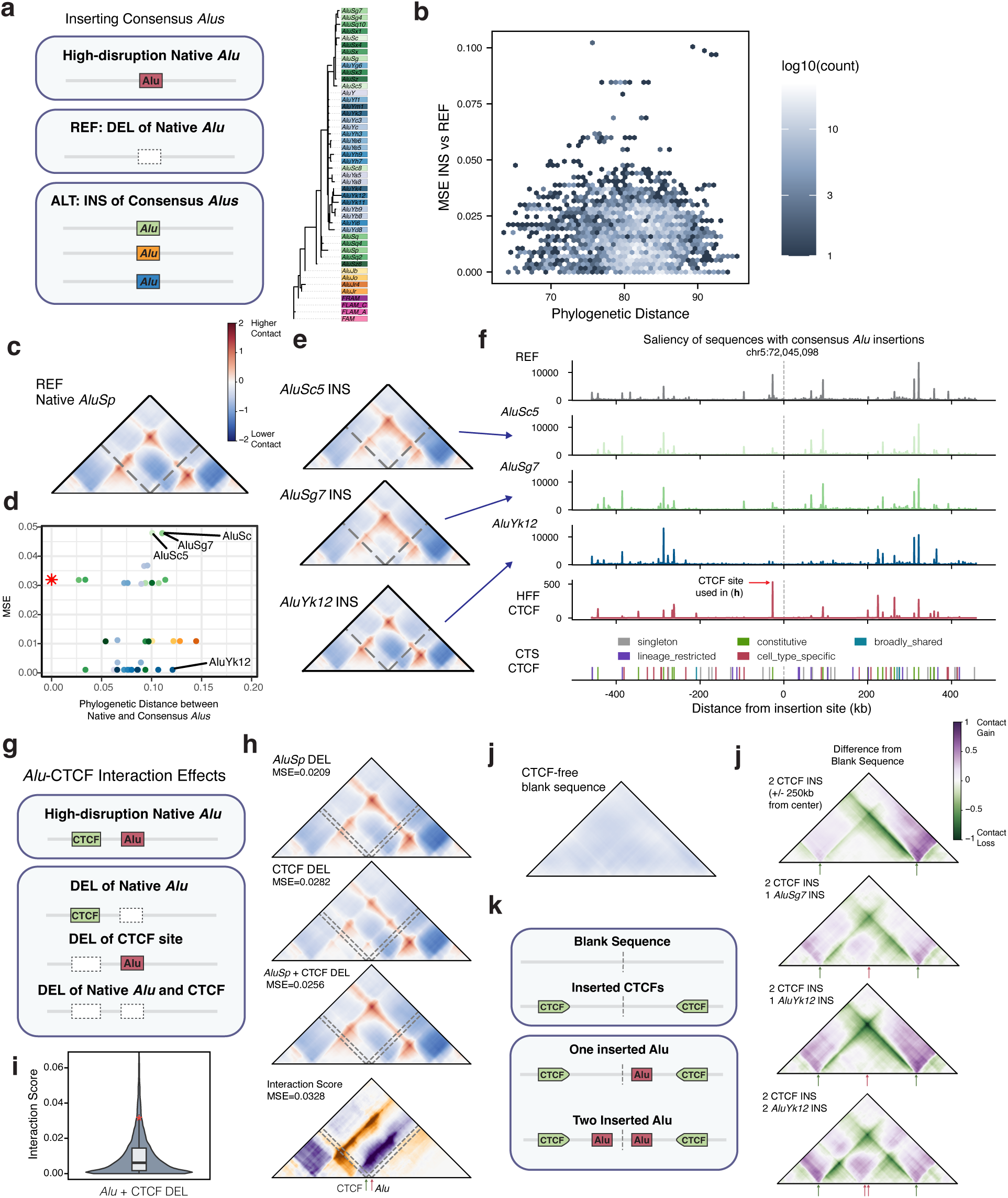
*Alu* elements differentially alter predicted chromatin folding by modulating CTCF boundary activity. **a.** Schematic of consensus *Alu* sequence insertion at loci containing high disruption *Alus*. **b.** Phylogenetic distance between native and inserted *Alu* sequences versus MSE between their predicted contact maps (Spearman ρ = 0.03). **c**. Predicted contact map for a native *AluSp* at chr5:72044951–72045246. **d.** MSE between the predicted contact maps of the native *AluSp* and each inserted consensus *Alu,* versus phylogenetic distance from the native *AluSp*. **e.** Predicted contact maps following insertion of *AluSc5*, *AluSg7*, and *AluYk12* consensus sequences at the locus shown in (**c**). **f.** Input saliency for the sequences containing the inserted *Alu* sequences shown in (**d**). **g.** Schematic of perturbations applied to the native *Alu* and nearby CTCF sites. **h.** Predicted contact maps following deletion of the native *Alu*, nearby CTCF sites, or both the *Alu* and CTCF site at the locus shown in (**c**). **i.** Interaction scores for *Alu*–CTCF deletion pairs. The star indicates the interaction score for the deletion shown in (**h**). **j.** Predicted contact map for a synthetic sequence lacking endogenous CTCF sites. **k.** Schematic illustrating the introduction of two inward-facing CTCF motifs into the CTCF-free sequence to generate a synthetic boundary, followed by insertion of one or two consensus *Alu* sequences. **l.** Predicted contact-map difference plots following insertion of one *AluSg7*, one *AluYk12*, or two *AluYk12* elements at the center of the synthetic CTCF boundary. Difference maps show changes relative to the sequence containing the two CTCF motifs alone.

To identify features that underlie these effects, we examined the Akita model’s saliency across the input sequences to measure the contribution of individual sequence positions to the model’s prediction (**Figure 6f, Methods**). Saliency at the inserted *Alu* itself remained low, whereas the strongest saliency peaks occurred at broadly occupied CTCF sites. Saliency at both neighboring and distal CTCF sites shifted depending on which *Alu* sequence was inserted, pointing to a sequence-specific interaction between *Alu* identity and CTCF binding.

We therefore hypothesized that Akita has learned long-range dependencies between *Alus* and CTCF sites. To test this, we further perturbed the *AluSp* locus in **Figure 6c**, deleting the CTCF site immediately to the left of the *Alu* that showed the strongest changes in saliency (**Figure 6f-g**). We compared deletion of (1) the native *Alu*, (2) the CTCF site alone, and (3) both the *Alu* and CTCF site. In this case, deleting the CTCF site alone introduced a new loop, whereas deleting both the *Alu* and CTCF site abolished this loop, suggesting a non-additive effect between the two elements (**Figure 6h**). To assess whether such interactions were specific to particular genomic pairs, we calculated interaction scores for paired deletions of *Alu–Alu*, *Alu– CTCF*, and *Alu–*random pairs across 1,000 randomly sampled highly disruptive *Alus* (**Supplementary Figure 15a, Supplementary Table 6, Methods**). The *Alu*–CTCF deletion at the locus in **Figure 6c** had an interaction score of 0.0328, larger in magnitude than those observed for randomly selected disruptive *Alu*–CTCF pairs (**Figure 6h-i, Supplementary Figure 15c,** *p* = 0.056). Two additional predicted CTCF perturbations at a different *AluY* locus showed similar effects (**Supplementary Figure 15a-b**), providing further evidence that *Alus* can interact non-additively with nearby CTCF sites.

Finally, to isolate *Alu*–CTCF interactions from genomic context, we performed a gain-of-function *in silico* experiment. First, we built a synthetic sequence containing no endogenous CTCF sites and introduced two inverted CTCF motifs 500kb apart. Akita predicted a weak TAD boundary from this minimal CTCF pair (**Figure 6j-k**). Inserting different *Alu* consensus sequences adjacent to the boundary altered the predicted folding pattern: for example, insertion of an *AluYk12* in the center between two CTCF sites strengthened the boundary, while inserting an *AluSg7* weakened the boundaries increased contacts elsewhere within the TAD (**Figure 6l**).

These effects also depended on the number of *Alus* present, with two inserted *AlusYk12* generating a new boundary near the insertion sites. Because these effects occurred in an otherwise featureless sequence, they demonstrate that *Alu* sequence variation can directly shape CTCF-dependent genome folding, providing a mechanistic basis for diverse, sequence-specific effects of *Alus* on genome folding.

## Discussion

Here, we used a machine learning framework to exhaustively evaluate the predicted effects of over 1 million *Alus* in the human genome on 3D genome folding. Our analyses suggest that *Alu*-mediated genome folding disruption is determined by two complementary factors. First, genomic context is broadly predictive: *Alus* in gene-rich, transcriptionally active regions are more likely to affect genome folding. Second, *Alu* sequence contributes additional disruptive potential. Even when genomic context is held constant, different *Alu* sequences produce distinct outcomes, ranging from strengthening existing CTCF-anchored boundaries to generating new loops or reshaping broader contact patterns. Thus, these results indicate that *Alus* do not influence genome architecture through a single uniform mechanism but instead exhibit a spectrum of effects shaped by both insertion site and their intrinsic sequence composition.

One of our most striking observations was that changing *Alu* sequence primarily redistributed model saliency across neighboring and distal CTCF sites rather than within the inserted *Alu* itself, as has been previously described ^45,46^. Given our synthetic perturbation experiments, this suggests that *Alus* may not act as independent architectural elements but instead modulate how existing CTCF sites contribute to boundary formation. *Alu* insertions may reweight regulatory circuitry to strengthen, weaken, or redirect CTCF-mediated interactions without necessarily introducing new binding sites for architectural proteins. More broadly, sequence determinants of genome folding extend beyond individual transcription factor motifs and are instead encoded through interactions among multiple sequence features distributed across the surrounding genomic landscape that likely influence chromatin states and conformations^47^.

This diversity of effects is consistent with a broader view of *Alus* as a source of evolutionarily innovation rather than uniformly deleterious mutations. Previous studies have shown that TE-derived sequences, including *Alus*, have been co-opted into new regulatory functions^8,13^. This regulatory re-wiring can involve local epigenetic changes, as exemplified by a heterozygous *AluY* insertion that generated expression of a novel MYBL2 isoform via loss of intronic CpG methylation ^48^. *Alu*-containing RNAs can also pair with one another through sequence complementarity and, in turn, target their homotypic *Alu* DNA loci — a mechanism shown to drive enhancer–promoter selectivity and organize transcribed regions around nuclear speckles ^12,39^, which was detectable using an Akita style model trained on RNA-DNA interactions ^49^.

Because *Alus* remain active and are polymorphic within human populations, these effects may contribute to inter-individual variation in gene expression via variable genome architecture ^50^, and, as a primate-specific element with ongoing lineage-specific activity, to inter-species divergence in genome architecture ^51–53^.

The molecular basis of these sequence-dependent effects merits further investigation. KZFPs recognize *Alu* elements and recruit chromatin-modifying machinery to establish heterochromatin, traditionally framed as a TE-KZFP arms-race^23,31,32^ However, KZFPs and TEs appear instead to cooperate, with KZFP control enabling rather than opposing TE co-option into transcriptional networks^37,42,54^. Two observations from our data support this model. First, disruptive *Alus* were associated with H3K27me3 but not H3K9me3, suggesting a more dynamic chromatin environment than the constitutive heterochromatin typical of KZFP-mediated silencing. Second, Akita assigns particularly high disruption scores to many *Alus* near genes involved in TE regulation, including KZFPs. We speculate this may reflect a self-limiting constraint: since these genes execute genome-wide TE silencing, silencing a nearby *Alu* could compromise their own expression and weaken silencing elsewhere in the genome. This offers a plausible reason for the persistence of disruptive, unsilenced *Alus*, and further support for a cooperative rather than antagonistic KZFP-TE relationship.

Our study has several limitations. First, all results depend on the Akita model, which captures one learned sequence grammar underlying genome folding. Models incorporating epigenomic information may capture additional context-specific determinants of genome folding not found in our analyses ^55^. However, these approaches require regulatory inputs beyond DNA sequence, making them difficult to apply to *in silico* sequence perturbations, where the appropriate epigenomic state of the altered locus is inherently unknown. Second, although we scored *Alu*s in both the hg38 and CHM13 T2T reference genomes, we used hg38 for most analyses because regulatory annotations and functional genomic datasets remain more extensively characterized in this assembly. As additional regulatory maps become available for T2T pangenomes, future studies will benefit from improved representation of repetitive and low-mappability regions. Finally, we must emphasize that our *in silico* predictions require experimental validation in the laboratory. However, the scale of this study would be infeasible with current experimental approaches, highlighting the utility of computational screening for generating mechanistic hypotheses and prioritizing loci for follow-up studies.

Our findings identify *Alus* as sequence elements that can shape three-dimensional genome organization through an interplay between their intrinsic sequence features and genomic context. Rather than having uniform effects, individual *Alus* span a spectrum of impacts on genome folding, depending on both their sequence and the regulatory landscapes in which they occur. As ongoing *Alu* retrotransposition continues today, these elements provide a persistent source of innovation, with the potential to reshape chromosome biology and gene regulation across multiple evolutionary timescales.

## Methods

### Data used

All data used in this study is publicly available. Analyses were performed in hg38 except for those involving 1000 Genomes data, where variants and tracks were aligned to the CHM13 T2T assembly.

For hg38, Repeatmasker, PhyloP, PhastCons, and mappability tracks were obtained from UCSC genome browser (https://hgdownload.soe.ucsc.edu/gbdb/hg38/). Gene expression, chromatin ChIP-seq tracks, whole genome bisulfite sequencing, and blacklist regions were obtained from ENCODE (encodeproject.org) for primary cells, cell lines, and in vitro differentiated cells. Gene annotations were from GENCODE v29. ChromHMM states were obtained from The Roadmap Epigenomics Project’s 15-state core model (https://egg2.wustl.edu/roadmap/web_portal/chr_state_learning.html). A curated panel of 60 epigenomes was used, spanning embryonic stem cells, induced pluripotent stem cells, primary cells, primary tissues, and several cell lines, many of which are shared with ENCODE.

For CHM13, SVs and TE annotations from long-read sequenced individuals were obtained from the 1000 Genomes Project ^29^. The CHM13 reference genome and Repeat masker annotations were obtained from the UCSC genome browser (https://hgdownload.soe.ucsc.edu/gbdb/hs1/), and gene annotations were obtained from the CHM13 genome repository (https://github.com/marbl/CHM13?tab=readme-ov-file).

### Scoring *Alus* with SuPreMo-Akita

We used the SuPreMo-Akita framework^20^ to score how disruptive an individual *Alu* is to genome organization. In total there were 1,181,071 *Alus* in the hg38 genome, but *Alus* that fell too close to centromeres and telomeres or *Alus* in windows with greater than 5% NA nucleotides were omitted, leaving 1,151,885 scored *Alus*. We scored, but then excluded both *Alus* and prediction windows that contain greater than 5% overlap with ENCODE blacklist regions leaving a total of 1,109,106 *Alus* for the remaining analyses ^56^.

For each *Alu*, we predicted the effect of deletion on genome folding by comparing the unperturbed (REF) and perturbed (ALT) contact maps predicted in the HFF cell type for the 1Mb region centered on the *Alu*. We then calculated a disruption score, which measures the mean squared error (MSE) or Spearman correlation (represented as CORR, or 1-(Spearman ρ)) between the REF and ALT maps. To reduce model-related technical biases, we used a sequence augmentation approach. For each input sequence, we generated six predictions: the original and reverse-complement sequences, each shifted 1 bp in either direction. We averaged the scores across these six augmentations to obtain a final disruption score for each alteration. We used MSE for most analyses, although we confirmed major trends were consistent with CORR. Finally, we defined ’highly disruptive’ *Alus* as those in the top 1% of deletion MSE scores, and ’neutral’ *Alu*s as those with MSEs at or below the 50th percentile.

For *Alus* and other SVs in the 1000 Genomes cohort, we took a similar approach to scoring, using the CHM13 T2T genome assembly. We score all biallelic insertions (INS) and deletions (DEL) annotated as *Alus* in the cohort. Additionally, we scored all biallelic SVs not annotated as a TE. We kept *Alus* and SVs between 275 to 375 bp, mirroring the distribution of *Alu* lengths in the cohort (median *Alu* length is 325 bp). After removing *Alus* and prediction windows overlapping blacklist regions ^56^, we were left with 18,137 and 1,859 *Alu* insertions and deletions, respectively, and 3,588 and 3,362 non-TE insertions and deletions, respectively.

### Assessing the relationship between *Alu* disruption and genomic sequence features

To assess the association between *Alu* disruption and different genomic features and characteristics, we looked at the intersection between each feature and the genomic region surrounding each *Alu*. Unless otherwise specified, genomic features were derived from HFF cells, matching the cell type used for SuPreMo-Akita predictions. DNA methylation was the exception, for which IMR90 whole-genome bisulfite sequencing (WGBS) data were used because IMR90 is a fibroblast cell type and therefore provides a closely related cellular context to HFF. To evaluate the contribution of cellular context, we repeated the analysis using genomic annotations from H1ESCs, while retaining the SuPreMo-Akita predictions generated in HFF.

We considered three types of genomic features, shown below:

| Feature type | Feature | Data reported |
| --- | --- | --- |
| Continuous signal | Mappability, phastCon100, GC content | Mean value per region |
| High frequency, length-varied, feature coverage | Repeatmasker, H3K9me3, H3K27me3, GENCODE genes, segmental duplications, self-chain regions | Fraction of overlap per region |
| Low-frequency, same-length features | Gene promoters, CTCF, *Alus*, WGBS, Accelerated and conserved sites derived from phyloP100 | Counts per region |

For continuous signals and GC content, we calculated the mean value over the region of interest surrounding each *Alu*. For coverage-based features, we calculated the proportion of overlap between each region and the union of all feature peaks, yielding values from zero to one. For count-based features, we calculated the number of elements falling within the region of interest. PhyloP100 scores >2 were classified as conserved sites (*phyloP100_high*), while scores <-2 were classified as accelerated sites (*phyloP100_low*). Each *Alu* was annotated at five window sizes: the *Alu* sequence itself, and the 1kb, 10kb, 100kb, and 1Mb regions surrounding the *Alu*. The 1Mb window corresponds to the Akita prediction window for that *Alu*.

To determine whether these genomic features were predictive of *Alu* disruption, we used two regression-based approaches. First, we used penalized logistic regression to distinguish highly disruptive *Alus* from neutral *Alus*. Second, we used elastic net regression to predict continuous disruption scores across all *Alus*. In both cases, we trained two models at each window size: one using the individual features, and one incorporating both the individual features and the pairwise interaction terms. Models were trained on all chromosomes except chromosome 2, which served as the held-out test set. Hyperparameters were selected via grid search over alpha values ranging from log(-4) to log(1), and L1 ratios ranging from 0.1 to 0.9, using five-fold cross validation with folds split by chromosomes. The hyperparameter combination yielding the lowest mean MSE across folds was selected for the final model.

### Identifying cell-type specific genes

We used bulk RNA-seq data from the ENCODE project and pulled all samples falling under the biosample classifications of cell lines, primary cells, and in vitro differentiated cells. We opted to not include tissues to eliminate the confounding factor of multiple cell types existing within a single tissue sample.

We categorized genes based on GENCODE annotations, aggregated across broader groups (https://www.gencodegenes.org/pages/biotypes.html). We focused our analyses on protein-coding genes, lncRNAs, lincRNAs, and pseudogenes, which were performed separately across the different biosamples. We kept only genes where at least one sample had a TPM greater than 5. For each gene, we calculated a Shannon entropy score from the TPM across all samples. A higher entropy score corresponds to a gene that is expressed in fewer cell types, and thus more cell-type specific.

### Classifying cell-type specific CTCF peaks and ChromHMM states

We identified cell-type specific CTCF ChIP-seq peaks and chromHMM states in analogous ways. First, we calculated a universe of all peaks by merging peaks that overlapped each other. For CTCF peaks, we used CTCF ChIP-seq from ENCODE encompassing all cell lines, primary cells, and in vitro differentiated cells, and excluding disease-related or perturbed samples.

Peaks were retained if they had a signal value of >10, -log(qval) > 4, and were less than 1000 bp long. For the chromHMM states, we used the 60 epigenomes that were trained and predicted on in the 15-state model ^35^. We merged CTCF peaks that had at least 100 bp of overlap, and chromHMM peaks that had at least a state-specific distance equal to half the median peak length for that state (computed from an initial unmerged union of peaks). From the universe of merged peaks, we defined five classes of cell-type specific peaks: (1) constitutive (>80% of samples), (2) broadly shared (50-80%), (3) lineage restricted (10-50%) (4) cell-type specific (2-10%), and (5) singletons (1 sample).

## 300bp random deletions

To understand the extent of genomic context to *Alu* disruption, we compared *in silico Alu* deletions to length-matched random deletions in the surrounding region. We sampled 1,000 *Alus* from the set of highly disruptive and neutral *Alu* sets. For each *Alu*, we defined a 10kb and 100kb window centered around the *Alu*, and sampled random, length-matched deletions within these windows, excluding other *Alus* and CTCF sites. To control for gene context, random deletions were constrained to fall within a gene if the original *Alu* was genic. For each *Alu*, we drew ten random deletions and reported the mean disruption score across the ten samples.

### *Alu* swaps within genes

We performed *in silico* swaps between highly disruptive and neutral *Alus* located within the same gene. For each gene, we first centered the prediction window at each ‘neutral’ *Alu.* For each of these centered ‘neutral’ *Alus*, we then iteratively replaced it with either a highly disruptive *Alu* or another neutral *Alu* from the same gene.

For each replacement, we calculate two disruption scores: (1) the MSE between the deletion of the query *Alu* and the REF sequence (contains the query *Alu*), and (2) the MSE between the deletion of the query *Alu* and the sequence in which the query *Alu* was replaced with the new *Alu*. This effectively allowed us to treat each *Alu* as an insertion and compare the insertion of different *Alus* into the same context.

We repeated this process for all ‘highly disruptive *Alus*’, inserting other highly disruptive and neutral *Alus* into all highly disruptive loci. Additionally, we tested only full length *Alus*, removing any *Alus* that were less than 275 bp long.

### ISM of GC content and CpG sites in *Alus*

To perturb individual sequence features we mutated individual nucleotides within a region of interest. To increase GC content, we calculated the total number of GC nucleotides in the region, and randomly converted a percentage of A/T nucleotides to C or G, with equal probability. Similarly, to decrease GC content, we selected a percentage of G/C nucleotides to convert to A or T. For CpG sites, we calculated the total number of CpGs within the region and introduced CpG sites by identifying TpG and CpA dinucleotides (the expected products of deamination of methylated CpGs) and reverting them to CpG via T→C (in TpG) or A→G (in CpA) substitutions, at a rate of either 50% or 100% above the original CpG count. Candidate sites were selected at random and non-overlapping positions were enforced so that no two edits could interfere with one another.

We performed GC and CpG mutagenesis on 1,000 randomly selected high-disruption and low-disruption Alus, filtering Alus to be full length (≥250bp) GC contents between the 25th and 75th percentiles across the sampled sites. For the flanking region analysis, we instead retained Alus with GC content between the 50th and 95th percentiles across both the Alu and its flanks, using a higher percentile cutoff to ensure a sufficient number of elements were retained.

### Identifying enriched transcription factor motifs in disruptive *Alus*

We identified overrepresented motifs in disruptive *Alus* using STREME and AME from the MEME suite ^57^. Known motifs were drawn from JASPAR 2026 and HOCOMOCO v14 ^58,59^. In both analyses, highly disruptive *Alus* served as the primary sequences and neutral *Alus* as the control sequences. We restricted sequences to 250 bp or greater to capture full-length *Alus* and avoid spurious motif discovery from truncated elements. Analyses were stratified by *Alu* family (*AluJ*, *AluS*, *AluY*).

### Rolling window analysis

To characterize positional effects of sequence disruption within *Alu* elements, we selected the 100 elements with the highest disruption score from each *Alu* subfamily. For each element, we deleted 30-bp windows tiled across its full genomic span at 10-bp intervals. For each deletion, we predicted the resulting contact map and calculated the disruption score as the MSE between the predicted contact map for the deleted sequence and that of the corresponding reference sequence. To aggregate scores across elements within each subfamily, we globally aligned each element’s sequence to its subfamily consensus, generating a per-element mapping from genomic to consensus coordinates. Each deletion window was then projected onto the corresponding consensus interval using this mapping. Scores were averaged across all elements at each consensus position to produce a subfamily-level profile of mean disruption score as a function of position along the *Alu* consensus sequence.

### *Alu* insertion analyses

To test the effect of a comprehensive range of *Alu* sequences on genome folding, we insert the 47 available consensus *Alu* sequences from the dfam database^44^ into the sites of highly disruptive *Alus*. We use the same random sampling of 1000 *Alus* from the set of ‘highly disruptive’ *Alus*. For each *Alu*, we center and delete the element. We then insert each consensus sequence in place of the native *Alu*, padding the sequence from the reference on either side. We calculated disruption from the predicted contact maps between the reference sequence containing the native *Alu* and the sequence with the consensus *Alu* inserted.

### Calculating evolutionary distances between *Alus*

Phylogenetic trees for *Alu* consensus sequences were constructed with the ‘msa’ package in R. First, the sequences were aligned with MUSCLE and a maximum-likelihood tree was built from the alignment, where a starting neighbor-joining tree was constructed using TN93 pairwise distances, and the best substitution model was selected by the optimal BIC. The tree was rooted on the ancestral FAM outgroup, and the final pairwise phylogenetic distances represent the summed branch lengths along the tree path between each pair of tips.

### Dual region deletions

To predict the effects of *Alus* interacting with *nearby* elements to alter genome folding, we performed *in silico* deletions of highly disruptive *Alus* together with a second genomic element. We randomly sampled 1000 *Alus* from the set of ‘highly disruptive’ *Alus*. For each sampled *Alu*, we identified candidate secondary elements within a 500 kb window centered on the *Alu* and, for each of four element types, sampled up to 50 secondary elements per original *Alu*: (1) another *Alu*; (2) another highly disruptive *Alu*; (3) a CTCF binding site (JASPAR MA0139.1 motif matches); (4) a length-matched random genomic region excluding other *Alus* within the window. *Alu* and CTCF-site sampling used a fixed random seed (random state=0); random-region coordinates were drawn uniformly at random from the remaining allowed intervals.

For all conditions, sequences were centered at the original *Alu* and contact map predictions were made for: (1) the REF sequence (no deletion) (2) original *Alu* deletion (3) deletion of the second element alone, and (4) the combined deletion of both the original *Alu* and secondary element. For cases involving deletion of the secondary element, we padded the input sequence from the reference genome on the side corresponding to the deleted region to maintain the 1Mb input length. Finally, we calculated the difference between the predicted contact maps for each perturbed sequence and the REF sequence.

To quantify interaction effects, we computed the signed, per-pixel deviation from additivity: for each pair, let δ₁, δ₂, and δ₁₂ denote the (ALT − REF) contact-map differences for the *Alu*-only, secondary-only, and combined deletions, respectively. The interaction score was defined as:

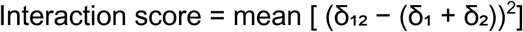

averaged over all valid (non-NaN) positions in the contact map. This measures whether the combined deletion’s effect matches the linear sum of the two elements’ individual effects: a score > 0 indicates the combined deletion is more disruptive than the sum of its parts (synergistic), while a score < 0 indicates it is less disruptive than expected (antagonistic).

### Calculating saliency contributions

Saliency was calculated for REF and consensus *Alu* insertion sequences. Per-nucleotide saliency was computed as the gradient of the summed Akita output with respect to the input sequence, via TensorFlow GradientTape. The four-channel (A/C/G/T) gradient at each position was then collapsed to a scalar by summing absolute values across channels. Per nucleotide saliency was aggregated into 2048 bp bins, totaling 448 bins per sequence, matching the Akita output. HFF CTCF ChIP-seq signal was binned over these same windows.

### *Alu* and CTCF insertions into a synthetic blank sequence

To test the sufficiency of *Alu* elements and CTCF motifs to reshape predicted 3D genome folding independent of native sequence context, we constructed a synthetic "blank" background sequence and used it as a canvas for controlled insertions. A 1 Mb sequence was generated by sampling nucleotides independently at each position, maintaining a GC content of 0.5. This sequence was then scanned against the JASPAR CTCF position weight matrix (MA0139.1), and any window scoring above a normalized threshold of 0.8 was iteratively replaced with newly sampled random sequence until no CTCF-like motifs remained.

To this background we introduced: (1) convergent CTCF motif pairs, using a CTCF "super motif" (TGGCCACTAGGTGGCGCCA) ^15^, placed symmetrically at varying distances (50–400 kb) from the sequence center and (2) *Alu* consensus sequences, inserted singly or in pairs again at symmetric distances from the center. We predicted contact maps for each of these sequences with Akita, and difference maps relative to the CTCF-free blank background and to CTCF-only controls were computed to isolate the folding contribution attributable specifically to the inserted *Alu* and/or CTCF elements.

### Statistical Tests

Statistical tests used include Pearson/Spearman correlations, Mann-Whitney U test, Fisher’s exact test, and the Kolmogorov-Smirnov test, either performed in R or Python.

## Data Availability

All data used in this study is publicly available.

## Code Availability

The SuPreMo-Akita code used to predict *Alu* disruption scores is located at https://github.com/ketringjoni/SuPreMo. Code for individual ISM experiments is forthcoming and will be available at https://github.com/shu-z/alu-3d-genome.

## Author Contributions

S.Z. and K.S.P. conceived this study. S.Z. performed computational analyses and wrote the initial draft. S.Z. and K.S.P. wrote the final manuscript.

## Acknowledgements

We thank the members of the Pollard lab for helpful discussions.

## References

1. Liao, X. et al. Repetitive DNA sequence detection and its role in the human genome. *Commun*. Biol. 6, 954 (2023).

2. Fudenberg, G. et al. Formation of chromosomal domains by loop extrusion. Cell Rep. 15, 2038–2049 (2016).

3. Rao, S. S. P. et al. Cohesin loss eliminates all loop domains. Cell 171, 305–320.e24 (2017).

4. Merkenschlager, M. & Nora, E. P. CTCF and cohesin in genome folding and transcriptional gene regulation. Annu. Rev. Genomics Hum. Genet. 17, 17–43 (2016).

5. Choudhary, M. N. K., Quaid, K., Xing, X., Schmidt, H. & Wang, T. Widespread contribution of transposable elements to the rewiring of mammalian 3D genomes. Nat. Commun. 14, 634 (2023).

6. Shukla, H., Huang, Y., Gao, Z. Y. & Lee, Y. C. G. Polymorphic 3D genome architecture mediated by transposable elements. bioRxivorg 2026.01.19.697799 (2026) doi:10.64898/2026.01.19.697799.

7. Huang, Y., Shukla, H. & Lee, Y. C. G. Species-specific chromatin landscape determines how transposable elements shape genome evolution. Elife 11, e81567 (2022).

8. Choudhary, M. N. et al. Co-opted transposons help perpetuate conserved higher-order chromosomal structures. Genome Biol. 21, 16 (2020).

9. Diehl, A. G., Ouyang, N. & Boyle, A. P. Transposable elements contribute to cell and species-specific chromatin looping and gene regulation in mammalian genomes. Nat. Commun. 11, 1796 (2020).

10. Batzer, M. A. & Deininger, P. L. Alu repeats and human genomic diversity. Nat. Rev. Genet. 3, 370–379 (2002).

11. Deininger, P. Alu elements: know the SINEs. Genome Biol. 12, 236 (2011).

12. Liang, L. et al. Complementary Alu sequences mediate enhancer-promoter selectivity. Nature 619, 868–875 (2023).

13. Chen, L.-L. & Yang, L. ALUternative regulation for gene expression. Trends Cell Biol. 27, 480–490 (2017).

14. Zhang, Y. et al. Transposable elements drive species-specific and tissue-specific transcriptomes in human development. Genome Biol. 26, 379 (2025).

15. Gunsalus, L. M., Keiser, M. J. & Pollard, K. S. In silico discovery of repetitive elements as key sequence determinants of 3D genome folding. Cell Genom. 3, 100410 (2023).

16. Zhou, J. Sequence-based modeling of three-dimensional genome architecture from kilobase to chromosome scale. Nat. Genet. 54, 725–734 (2022).

17. Tan, J. et al. Cell-type-specific prediction of 3D chromatin organization enables high-throughput in silico genetic screening. Nat. Biotechnol. 41, 1140–1150 (2023).

18. Avsec, Ž., et al. Advancing regulatory variant effect prediction with AlphaGenome. Nature 649, 1206–1218 (2026).

19. Chen, V. et al. Applying interpretable machine learning in computational biology-pitfalls, recommendations and opportunities for new developments. Nat. Methods 21, 1454–1461 (2024).

20. Gjoni, K. & Pollard, K. S. SuPreMo: a computational tool for streamlining in silico perturbation using sequence-based predictive models. Bioinformatics 40, btae340 (2024).

21. Fudenberg, G., Kelley, D. R. & Pollard, K. S. Predicting 3D genome folding from DNA sequence with Akita. Nat. Methods 17, 1111–1117 (2020).

22. Gjoni, K. et al. Comparing chromatin contact maps at scale: methods and insights. Nat. Methods 22, 824–833 (2025).

23. Davis, J., Voicu, D., Chitnavis, U., Jaksina, J. & Imbeault, M. The role of KRAB zinc-finger proteins in expanding the domestication potential of transposable elements. Nat. Genet. 58, 492–502 (2026).

24. Noonan, J. P., Grimwood, J., Schmutz, J., Dickson, M. & Myers, R. M. Gene conversion and the evolution of protocadherin gene cluster diversity. Genome Res. 14, 354–366 (2004).

25. Duan, R. V. et al. Widespread distribution of Alu/Alu-mediated genomic rearrangement predisposing to a broad range of Mendelian disease and cancer in human populations. Genome Med. 18, (2026).

26. Balachandran, P. et al. Transposable element-mediated rearrangements are prevalent in human genomes. Nat. Commun. 13, 7115 (2022).

27. Ferrari, R. et al. TFIIIC binding to Alu elements controls gene expression via chromatin looping and histone acetylation. Mol. Cell 77, 475–487.e11 (2020).

28. Zhang, X., Fang, B. & Huang, Y.-F. Transcription factor binding sites are frequently under accelerated evolution in primates. Nat. Commun. 14, 783 (2023).

29. Schloissnig, S. et al. Structural variation in 1,019 diverse humans based on long-read sequencing. Nature 644, 442–452 (2025).

30. Konkel, M. K. et al. Sequence analysis and characterization of active human Alu subfamilies based on the 1000 genomes pilot project. Genome Biol. Evol. 7, 2608–2622 (2015).

31. Zhou, W., Liang, G., Molloy, P. L. & Jones, P. A. DNA methylation enables transposable element-driven genome expansion. Proc. Natl. Acad. Sci. U. S. A. 117, 19359–19366 (2020).

32. Zhang, X.-O., Gingeras, T. R. & Weng, Z. Genome-wide analysis of polymerase III-transcribed Alu elements suggests cell-type-specific enhancer function. Genome Res. 29, 1402–1414 (2019).

33. Chuong, E. B., Elde, N. C. & Feschotte, C. Regulatory activities of transposable elements: from conflicts to benefits. Nat. Rev. Genet. 18, 71–86 (2017).

34. Walter, M., Teissandier, A., Pérez-Palacios, R. & Bourc’his, D. An epigenetic switch ensures transposon repression upon dynamic loss of DNA methylation in embryonic stem cells. Elife 5, e11418 (2016).

35. Ernst, J. & Kellis, M. ChromHMM: automating chromatin-state discovery and characterization. Nat. Methods 9, 215–216 (2012).

36. Zamudio, N. & Bourc’his, D. Transposable elements in the mammalian germline: a comfortable niche or a deadly trap? Heredity (Edinb.) 105, 92–104 (2010).

37. Pontis, J. et al. Hominoid-specific transposable elements and KZFPs facilitate human embryonic genome activation and control transcription in naive human ESCs. Cell Stem Cell 24, 724–735.e5 (2019).

38. Karczewski, K. J. et al. The mutational constraint spectrum quantified from variation in 141,456 humans. Nature 581, 434–443 (2020).

39. Liu, S. et al. Alu repeat-containing RNAs spatially organize actively transcribed genomic regions around nuclear speckles. Mol. Cell 86, 1723–1741.e11 (2026).

40. Radman-Livaja, M. & Rando, O. J. Nucleosome positioning: how is it established, and why does it matter? Dev. Biol. 339, 258–266 (2010).

41. Lorch, Y., Maier-Davis, B. & Kornberg, R. D. Role of DNA sequence in chromatin remodeling and the formation of nucleosome-free regions. Genes Dev. 28, 2492–2497 (2014).

42. Imbeault, M., Helleboid, P.-Y. & Trono, D. KRAB zinc-finger proteins contribute to the evolution of gene regulatory networks. Nature 543, 550–554 (2017).

43. Liao, T.-T. et al. Let-7 modulates chromatin configuration and target gene repression through regulation of the ARID3B complex. Cell Rep. 14, 520–533 (2016).

44. Storer, J., Hubley, R., Rosen, J., Wheeler, T. J. & Smit, A. F. The Dfam community resource of transposable element families, sequence models, and genome annotations. Mob. DNA 12, 2 (2021).

45. Zhu, K., Zhuo, J. & Zhen, Y. Evolution of CTCF binding sites in the human genome. Mol. Biol. Evol. 43, msag167 (2026).

46. Schmidt, D. et al. Waves of retrotransposon expansion remodel genome organization and CTCF binding in multiple mammalian lineages. Cell 148, 335–348 (2012).

47. Schwalie, P. C. et al. Co-binding by YY1 identifies the transcriptionally active, highly conserved set of CTCF-bound regions in primate genomes. Genome Biol. 14, R148 (2013).

48. Lucas, J. K., et al. HPRC2: A human pangenome reference with near-complete coverage of common genetic variation. bioRxivorg 2026.07.21.739710 (2026) doi:10.64898/2026.07.21.739710.

49. Kuang, S. & Pollard, K. S. Exploring the roles of RNAs in chromatin architecture using deep learning. Nat. Commun. 15, 6373 (2024).

50. Gilbertson, E. N. et al. Machine learning reveals the diversity of human 3D chromatin contact patterns. Mol. Biol. Evol. 41, msae209 (2024).

51. Brand, C. M. et al. Sequence-based machine learning reveals 3D genome differences between bonobos and chimpanzees. Genome Biol. Evol. 16, evae210 (2024).

52. Keough, K. C. et al. Three-dimensional genome rewiring in loci with human accelerated regions. Science 380, eabm1696 (2023).

53. Thybert, D. et al. Repeat associated mechanisms of genome evolution and function revealed by the Mus caroli and Mus pahari genomes. Genome Res. 28, 448–459 (2018).

54. Turelli, P. et al. Primate-restricted KRAB zinc finger proteins and target retrotransposons control gene expression in human neurons. Sci. Adv. 6, eaba3200 (2020).

55. Bai, J. et al. High-throughput in silico screen uncovers key regulators of 3D genome architecture. bioRxivorg 2025.12.09.693120 (2025) doi:10.64898/2025.12.09.693120.

56. Ogata, J. D. et al. excluderanges: exclusion sets for T2T-CHM13, GRCm39, and other genome assemblies. Bioinformatics 39, btad198 (2023).

57. Bailey, T. L., Johnson, J., Grant, C. E. & Noble, W. S. The MEME suite. Nucleic Acids Res. 43, W39–49 (2015).

58. Ovek Baydar, D., et al. JASPAR 2026: expansion of transcription factor binding profiles and integration of deep learning models. Nucleic Acids Res. 54, D184–D193 (2026).

59. Vorontsov, I. E. et al. HOCOMOCO in 2024: a rebuild of the curated collection of binding models for human and mouse transcription factors. Nucleic Acids Res. 52, D154–D163 (2024).

