## Supplementary Information for "Machine learning reveals sequence and genomic context features underlying *Alu*-specific effects on genome folding"

**Supplementary Figures** **2**
Supplementary Figure 1 2
Supplementary Figure 2 3
Supplementary Figure 3 4
Supplementary Figure 4 5
Supplementary Figure 5 6
Supplementary Figure 6 7
Supplementary Figure 7 8
Supplementary Figure 8 10
Supplementary Figure 9 11
Supplementary Figure 10 12
Supplementary Figure 11 13
Supplementary Figure 12 14
Supplementary Figure 13 15
Supplementary Figure 14 16
Supplementary Figure 15 17
**Supplementary Tables** **18**

### Supplementary Figures

#### Supplementary Figure 1

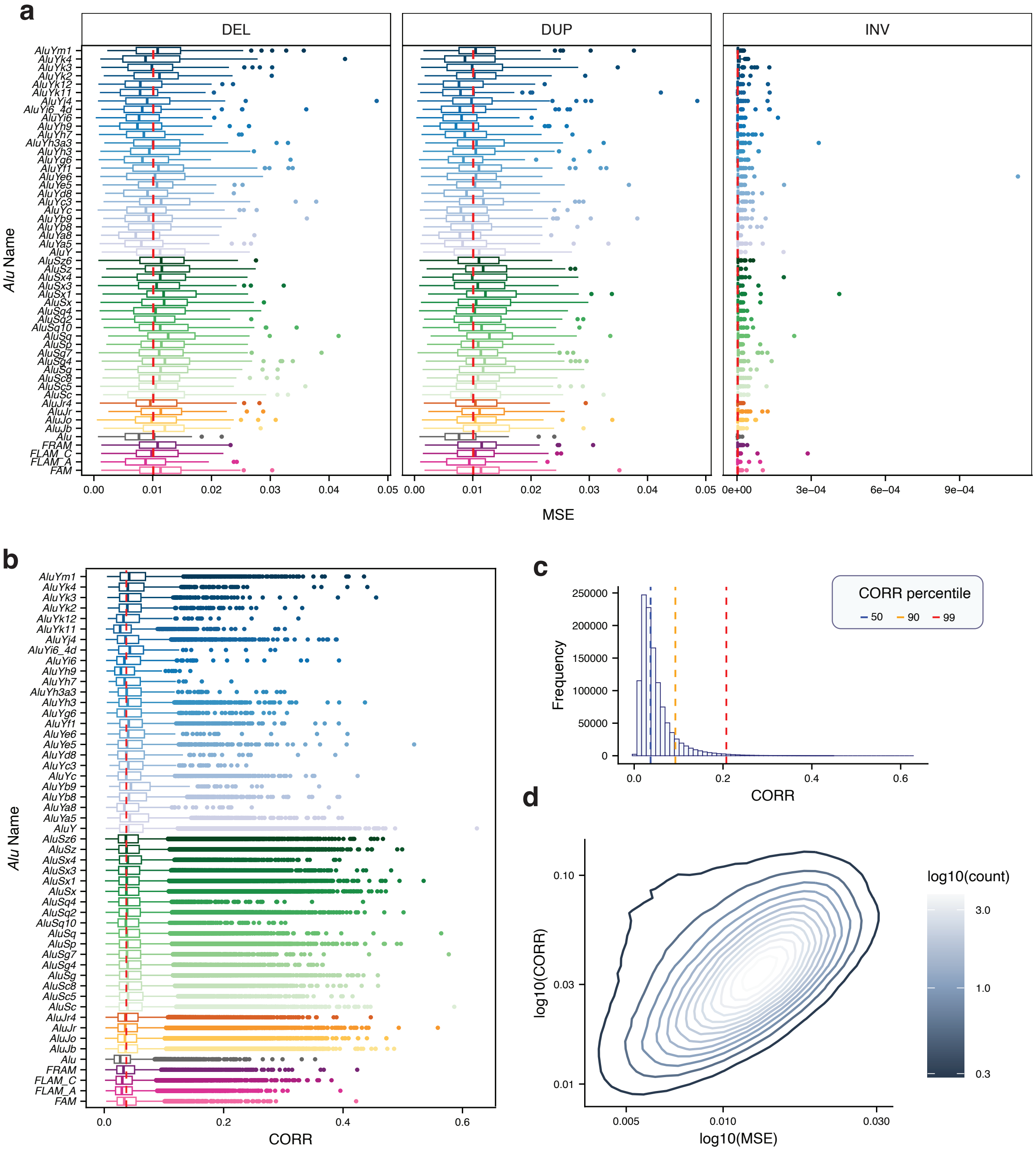


**Supplementary Figure 1. Additional disruption metrics for all *Alus* in hg38**. **a.** Disruption scores for deletions (mean MSE=0.010), duplications (mean MSE=0.010), and inversions (mean MSE $6.31 \times 10^{-6}$) of *Alus.* 100 *Alus* per *Alu* type are represented (n=5100). Deletions and duplication MSE scores were highly correlated (Pearson *R*=0.863). **b.** Distribution of CORR disruption scores across all *Alus*, grouped by *Alu* type. The dashed red line represents the median CORR score across all *Alus*. **c.** Distribution of CORR disruption scores. Dashed lines indicate CORR percentiles. **d.** Comparison of MSE and CORR score distributions.

### Supplementary Figure 2

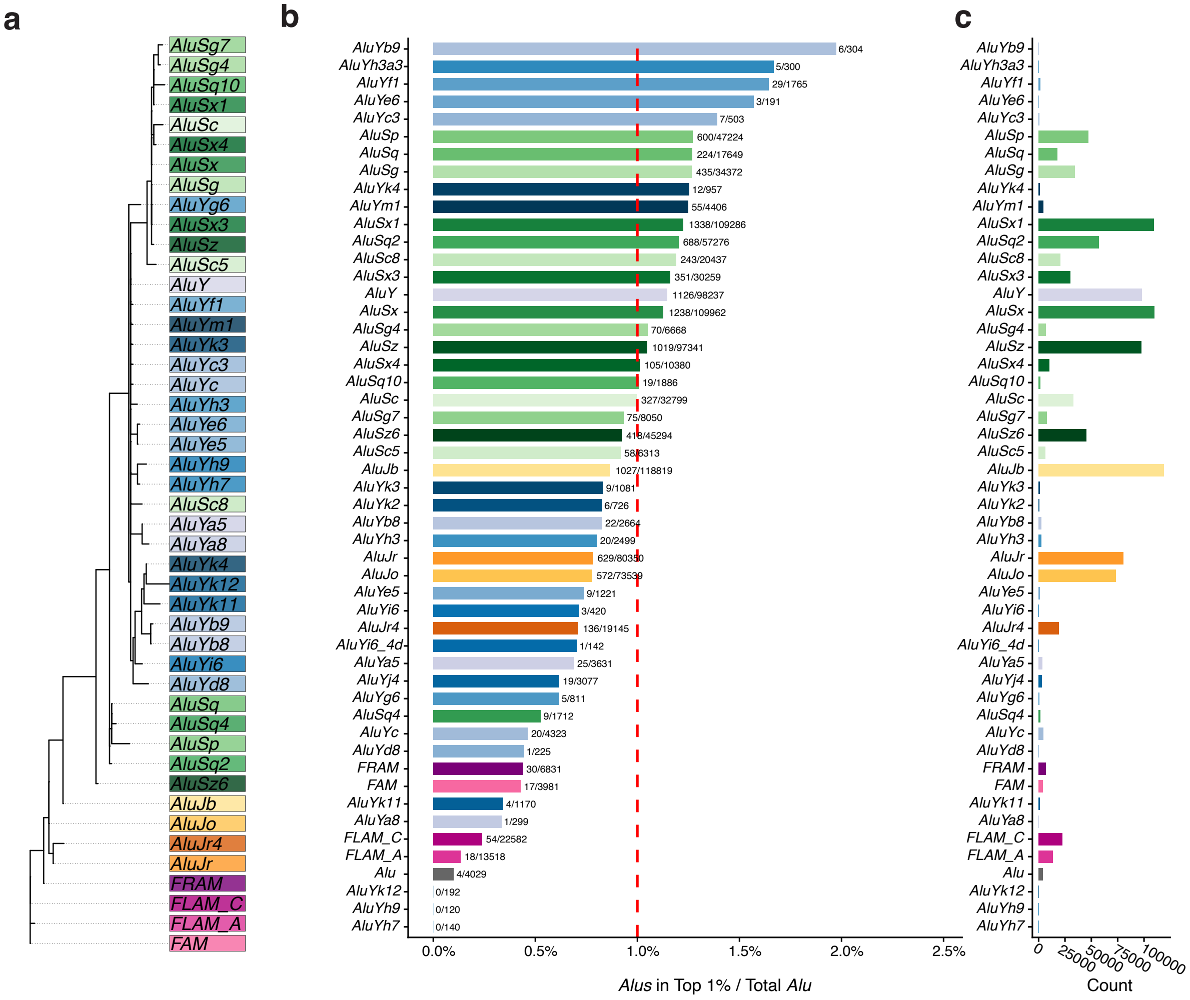


**Supplementary Figure 2. Younger *Alus* are enriched among disruptive *Alus*. a**. Phylogenetic tree of *Alu* consensus sequences, where branch lengths represent sequence divergence, used as a proxy for evolutionary age. *Alus* are colored by the broader *Alu* family they belong to. **b.** Proportion of *Alus* per *Alu* type that are highly disruptive (top 1% MSE). c. Total number of *Alus* in the genome, per *Alu* type.

### Supplementary Figure 3

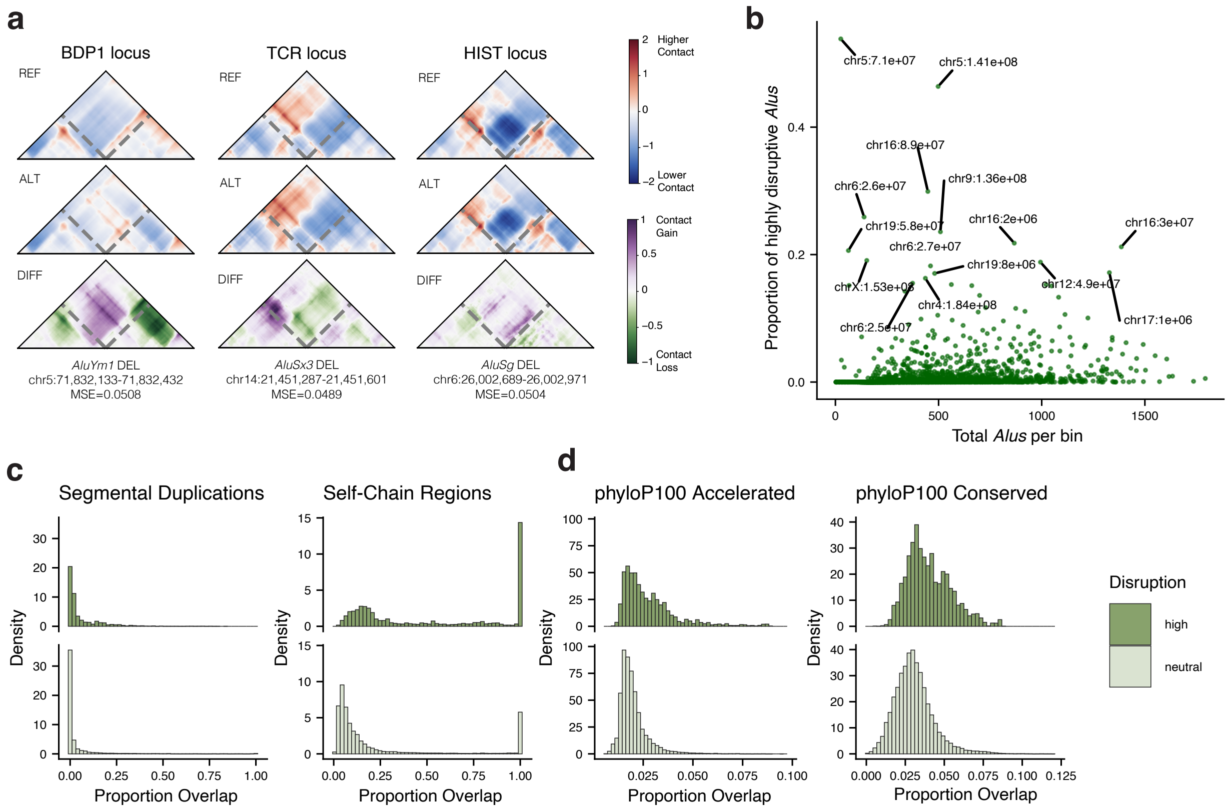


**Supplementary Figure 3. Disruptive *Alus* localize to regions with high evolutionary turnover. a.** Example maps of *Alus* located in 1Mb disruption hotspot bins, at the BDP1 locus, T-cell receptor gene cluster, and histone gene cluster. **b.** Relationship between total *Alu* density and disruptive *Alu* density, for 1Mb bins across the genome. **c-d.** Overlap distributions for segmental duplications, self-chain regions, regions of low and high sequence conservation, for highly disruptive versus neutral *Alus* (1 Mb window, Mann-Whitney U $p < 2.2\times10^{-16}$ for all comparisons).

### Supplementary Figure 4

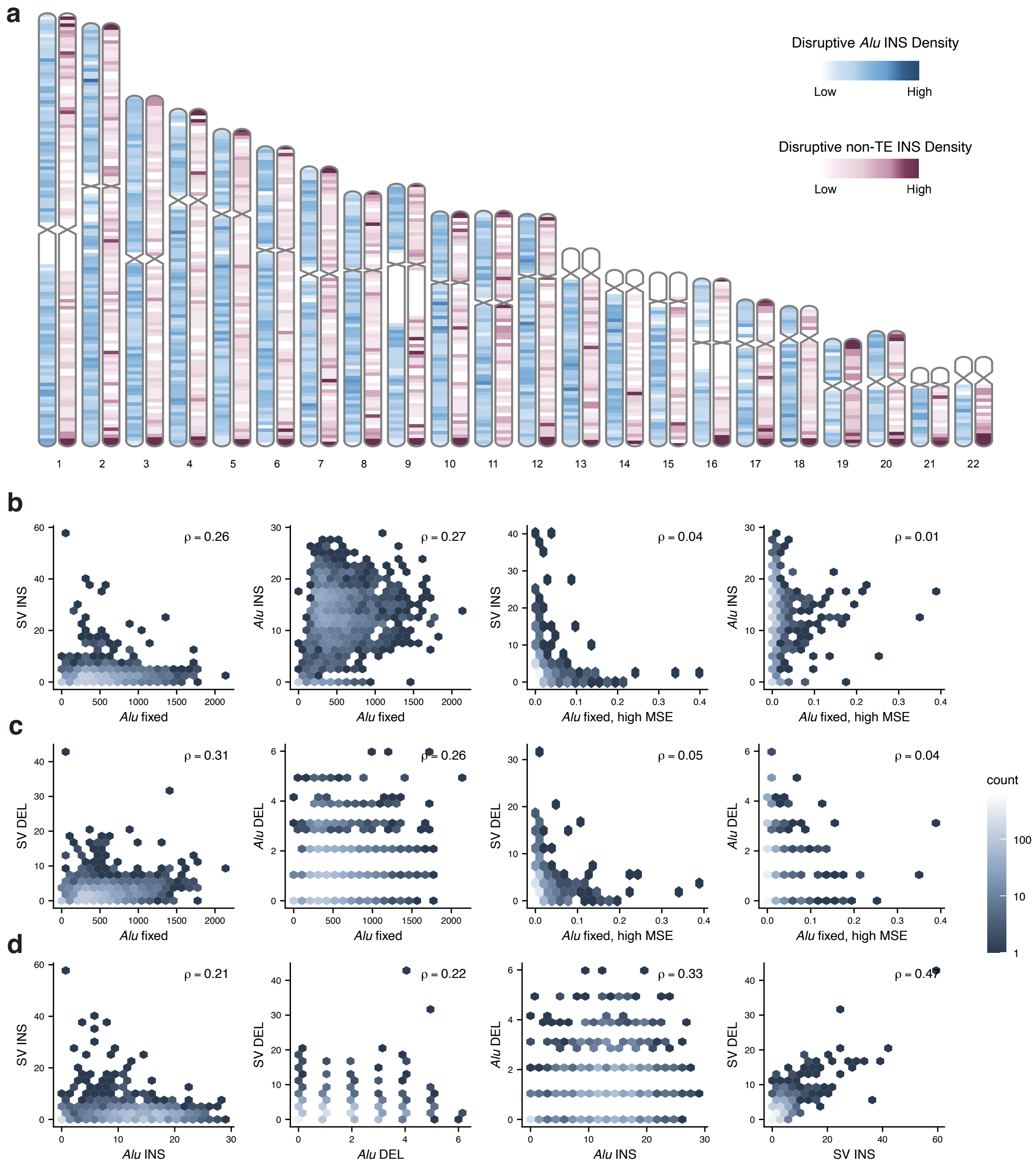


**Supplementary Figure 4. Polymorphic *Alus* show a distinct genomic distribution from non-TE SVs. a.** Genome-wide distribution of non-TE insertions and polymorphic *Alu* insertions, in 2 Mb bins. **b-c.** Genomic distribution of fixed *Alus* (all and highly disruptive) compared to polymorphic *Alu* and non-TE insertions (**b**) and deletions (**c**). **d.** Genome-wide relationship between non-TE and *Alu* insertion/deletion density. Spearman correlations are reported in b-d.

### Supplementary Figure 5

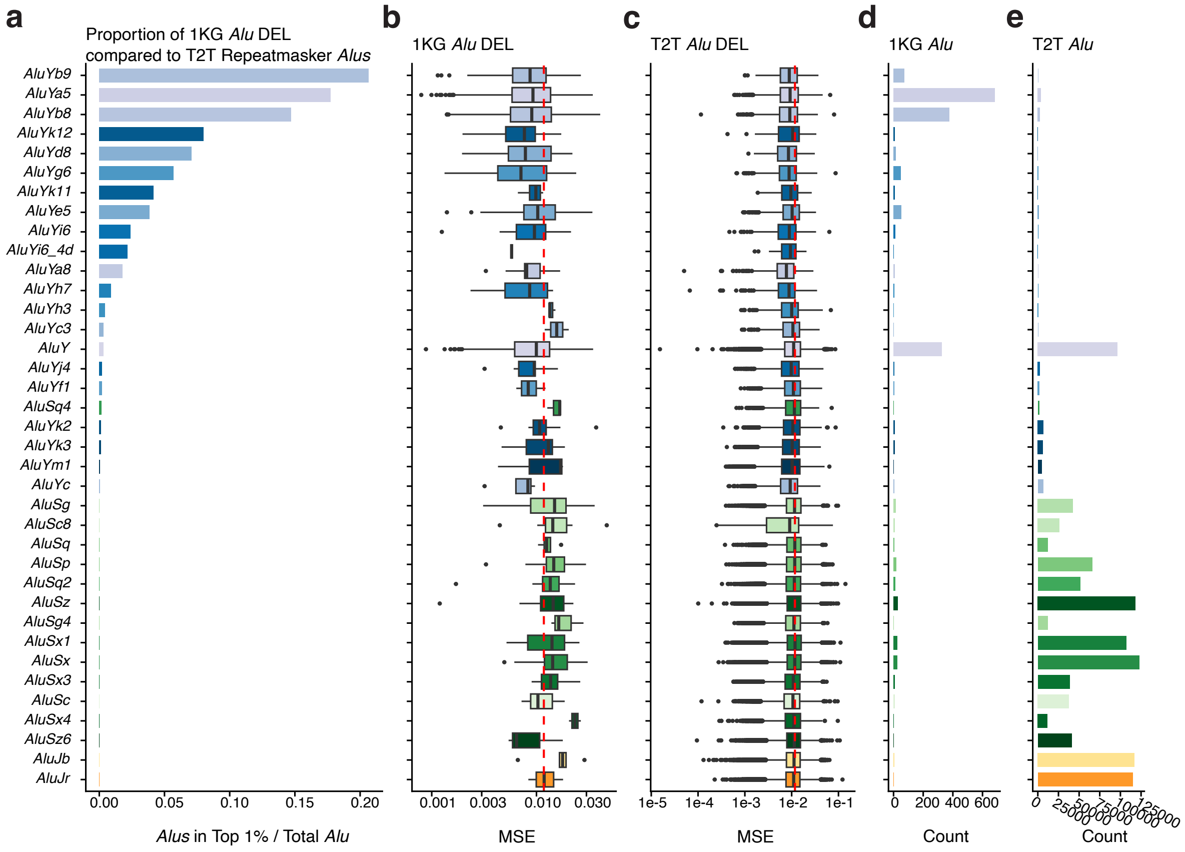


**Supplementary Figure 5. Younger *Alu* families disproportionately make up the set of highly disruptive polymorphic *Alu* deletions. a.** Proportion of polymorphic *Alu* deletions that are highly disruptive. **b.** Distribution of disruption scores for all polymorphic *Alu* deletions. The red dashed line indicates the 99th percentile of MSE disruption (MSE>0.0288) for CHM13 *Alus*. **c.** Distribution of disruption scores for all fixed *Alu* deletions in CHM13. **d.** The total number of polymorphic *Alu* deletions, sorted by *Alu* family. **e.** The total number of fixed *Alus* in CHM13, sorted by *Alu* family.

### Supplementary Figure 6

**a**

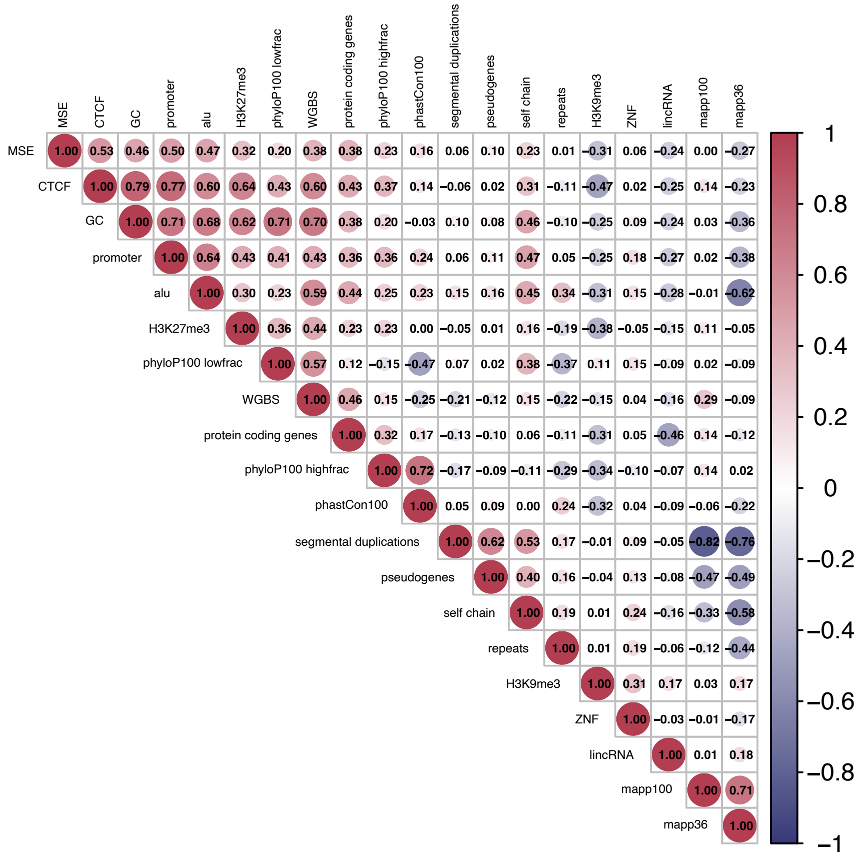


**Supplementary Figure 6. Genomic features of *Alus* tend to be correlated with each other.**
**a.** Pearson correlation of all context features used in the regression models to classify or predict disruptive *Alus*, calculated from the 1Mb Akita prediction window.

### Supplementary Figure 7

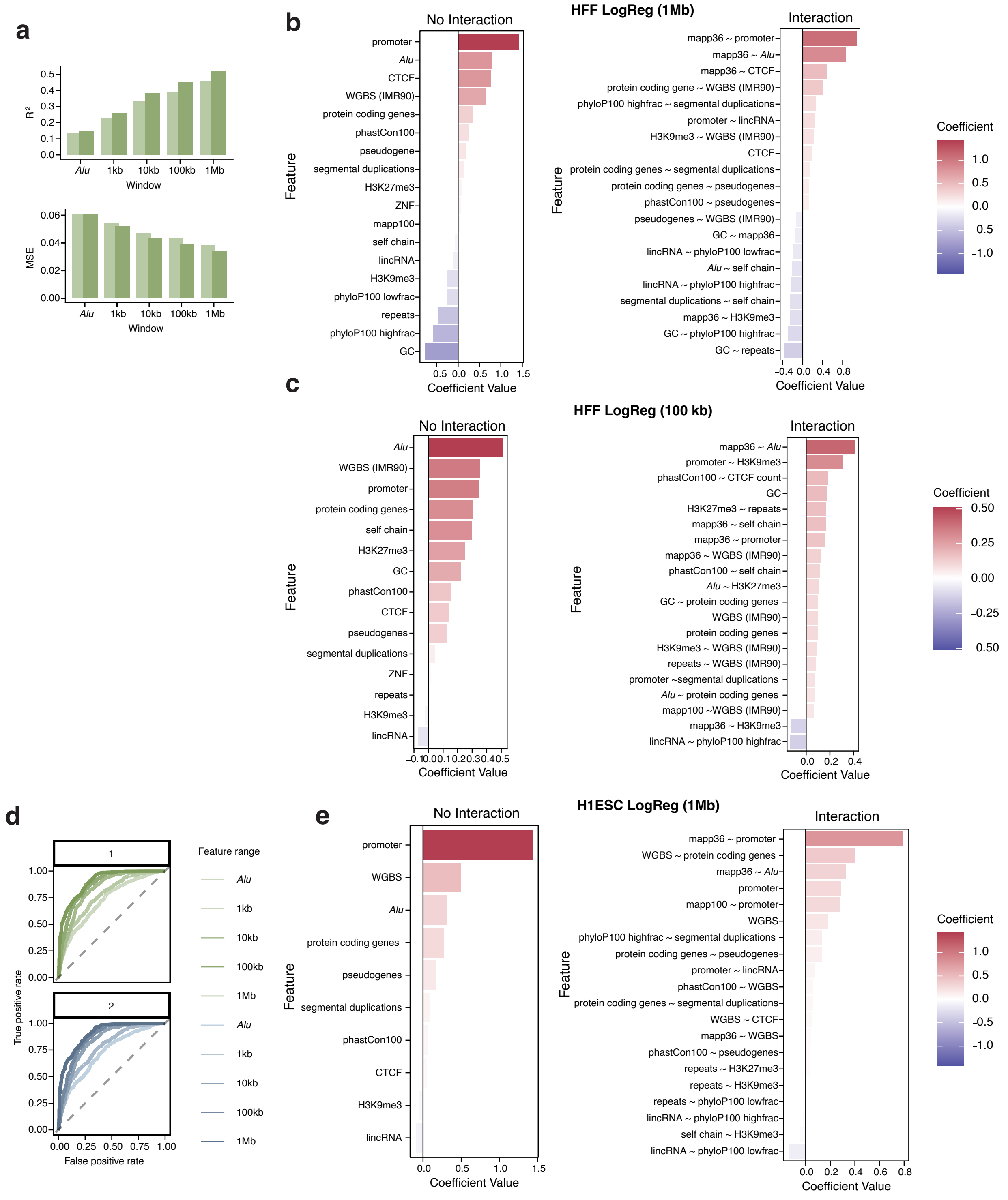

**Supplementary Figure 7. Predictive genomic features of Alu-associated genome folding disruption are largely consistent across regression models and cell types. a.** MSE and $R^2$ performance of elastic net models on predicting *Alu* disruption from features in the surrounding genomic window (HFF), ranging from the length of the *Alu* itself to the full 1 Mb Akita prediction window. Models were fit using individual features alone and with pairwise feature interactions **b.** Coefficients of the most predictive features for the no-interaction and interaction-term logistic regression models trained on the 1Mb window (HFF). **c.** Coefficients of the most predictive features from the logistic regression model trained on the 100 kb window (HFF). **d.** AUROC curves for the logistic regression models trained on the 1 Mb window (H1-ESC). **e.** Coefficients of the most predictive features for the no-interaction and interaction-term logistic regression models (H1ESC).

### Supplementary Figure 8

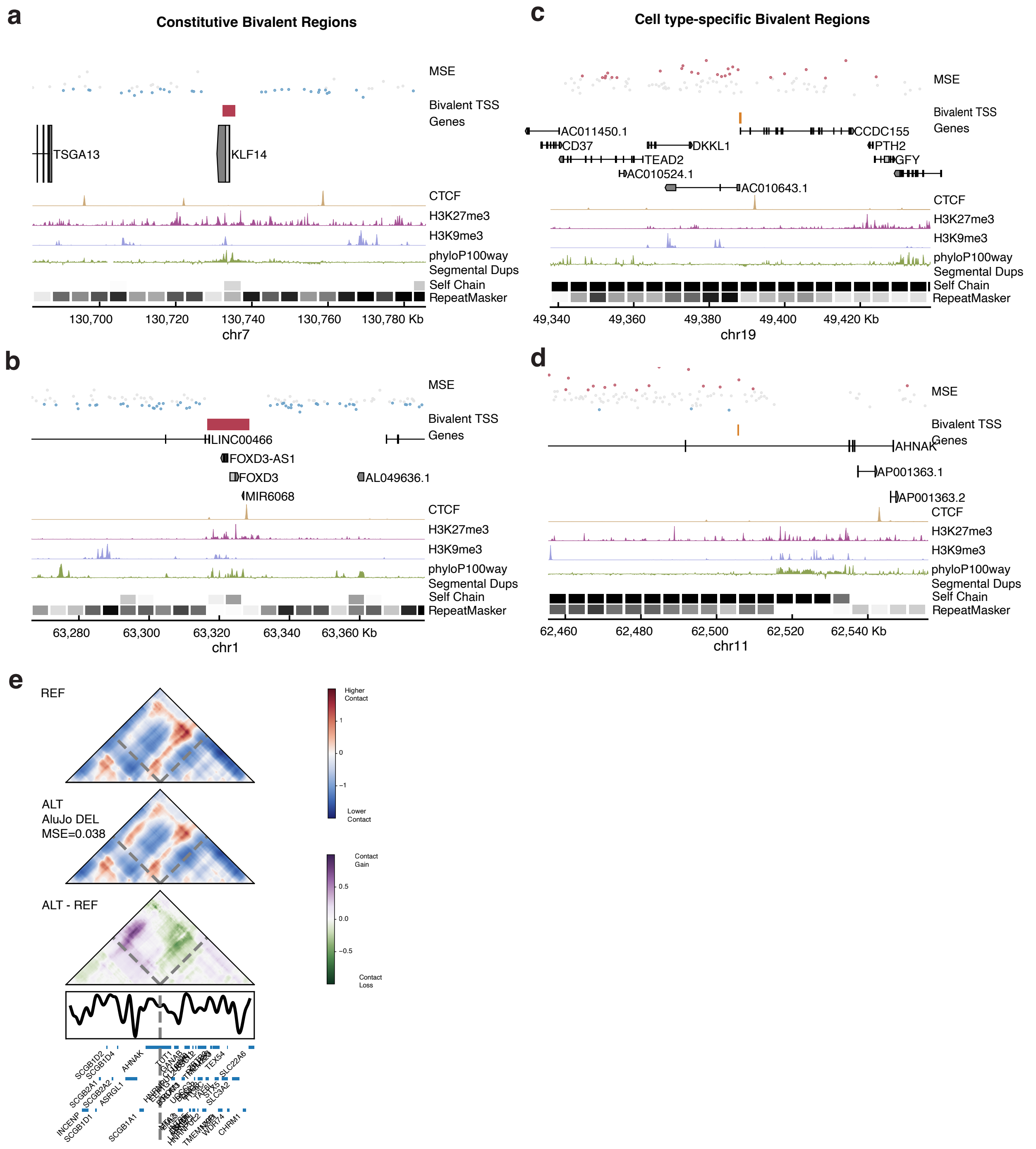


**Supplementary Figure 8. Cell type specific bivalent ChromHMM regions tend to be located near disruptive *Alus*. a-b.** Genome browser tracks of constitutive bivalent ChromHMM regions (State 10) on chr7 (**a**) and chr1 (**b**), with *Alu* elements marked in blue (neutral) or red (highly disruptive). **c-d.** Genome browser tracks of cell-type specific bivalent ChromHMM regions (State 10) on chr19 (**c**) and chr11 (**d**), with *Alus* colored as in (**a-b**). **e.** Predicted contact map for a deletion of a highly disruptive *AluY* from the cell-type specific bivalent region in (**d**).

### Supplementary Figure 9

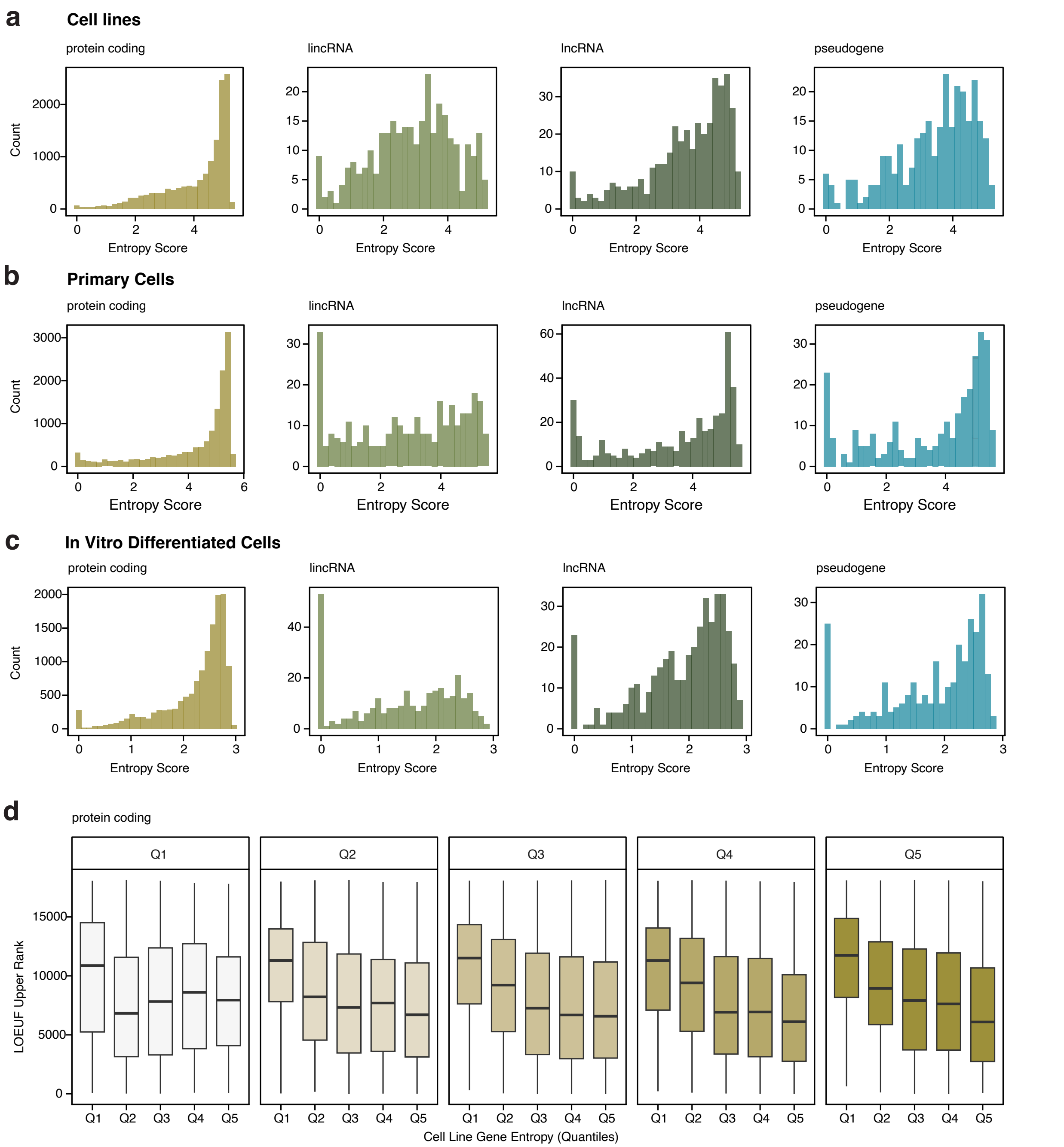


**Supplementary Figure 9. Genes with broader expression tend to harbor more disruptive *Alus*, which holds regardless of biosample type. a–c.** Entropy score distributions for protein-coding genes, lincRNAs, lncRNAs, and pseudogenes, computed within (**a**) cell lines, (**b**) primary cells, and (**c**) in vitro differentiated cells from the ENCODE consortium. **d.** LOEUF score (y-axis) versus entropy score (x-axis) for protein-coding genes, faceted by average *Alu* disruption MSE within each gene (quantiles). Genes with higher expression entropy and more disruptive *Alus* tend to have lower LOEUF scores, indicating greater evolutionary constraint.

### Supplementary Figure 10

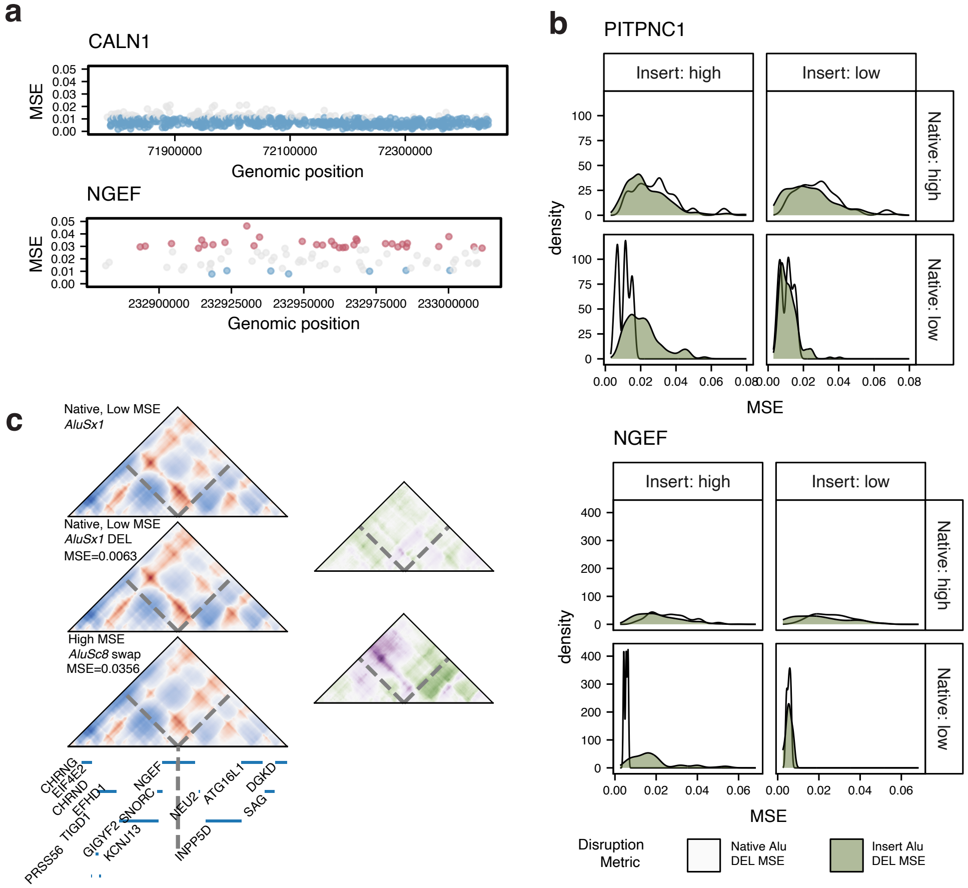


**Supplementary Figure 10. *Alu*-swap results in individual genes. a.** Distributions of *Alu* disruption within *CALN1*, a gene with high *Alu* density but broadly low disruption, and *NGEF*, a gene containing many disruptive *Alus* along with a wide range of disruption across the gene body. **b.** Predicted disruption after swapping high- and low-disruption *Alus* within the same gene, shown for all four native/insert combinations in *PITPNC1* and *NGEF*. Within each combination, the white shading reflects disruption from the native *Alu's* deletion, and while the green shading reflects the disruption introduced by the swapped-in *Alu*, relative to that native-deletion baseline. **c.** Predicted contact maps in NGEF upon deletion of a native low-MSE *AluSx1*, followed by insertion of a high-MSE *AluSc8* in its place.

### Supplementary Figure 11

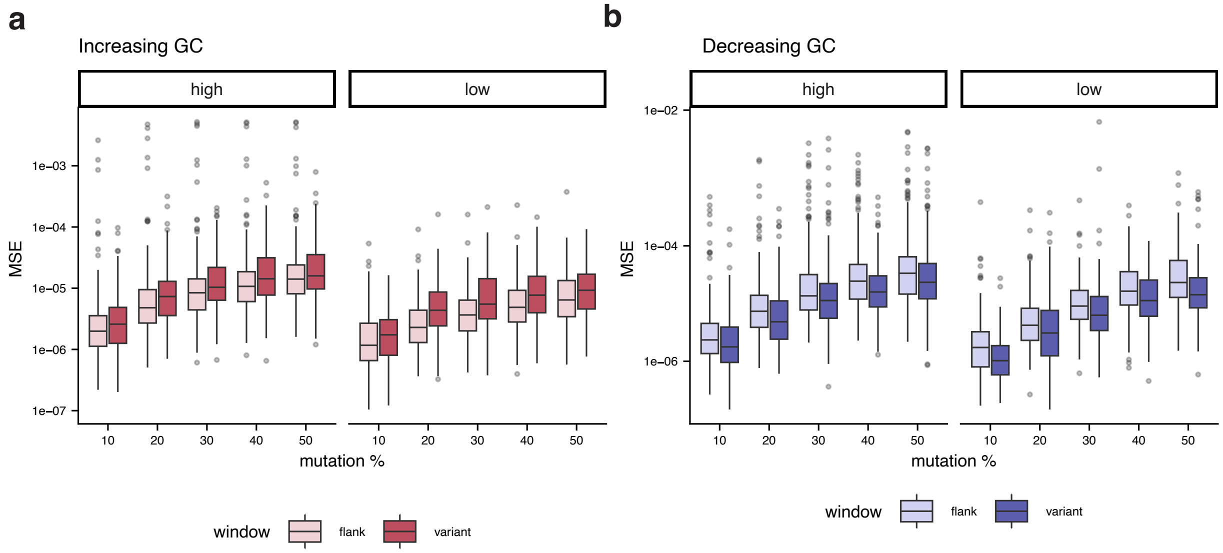


**Supplementary Figure 11. GC mutagenesis of *Alu* and flanking regions. a.** Increasing GC content in the *Alu* versus flanking regions. **b.** Decreasing GC content in the *Alu* versus flanking regions. The *Alu* and flanking regions were matched for initial GC content.

### Supplementary Figure 12

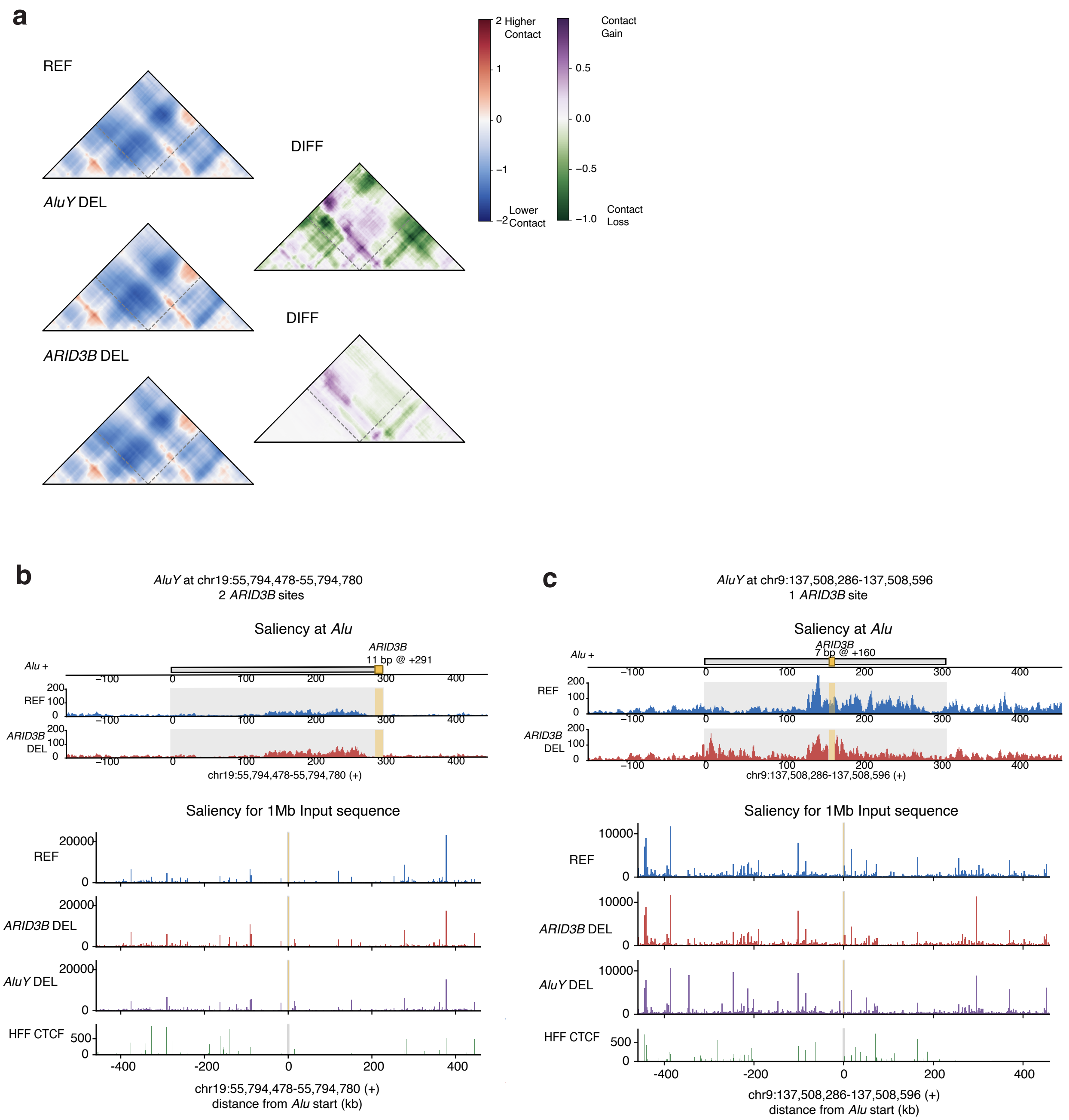


**Supplementary Figure 12. Model saliency for *ARID3B* motif perturbation within *AluY* elements. a.** Input gradients following deletion of an *ARID3B* motif at the end of an *AluY* element on chr19, shown at both the *Alu* window (base-pair resolution) and the Akita prediction window (2,048-bp resolution). The gradients highlight the contribution of sequence positions within the input sequence to the predicted genome-folding pattern. **b.** Input gradients for an *AluY* element on chr9 in which the *ARID3B* motif is located near the center of the element. **c.** Predicted contact maps for the region shown in (**b**) under three conditions: the unperturbed sequence, deletion of the entire *AluY* element, and deletion of the *ARID3B* motif.

### Supplementary Figure 13

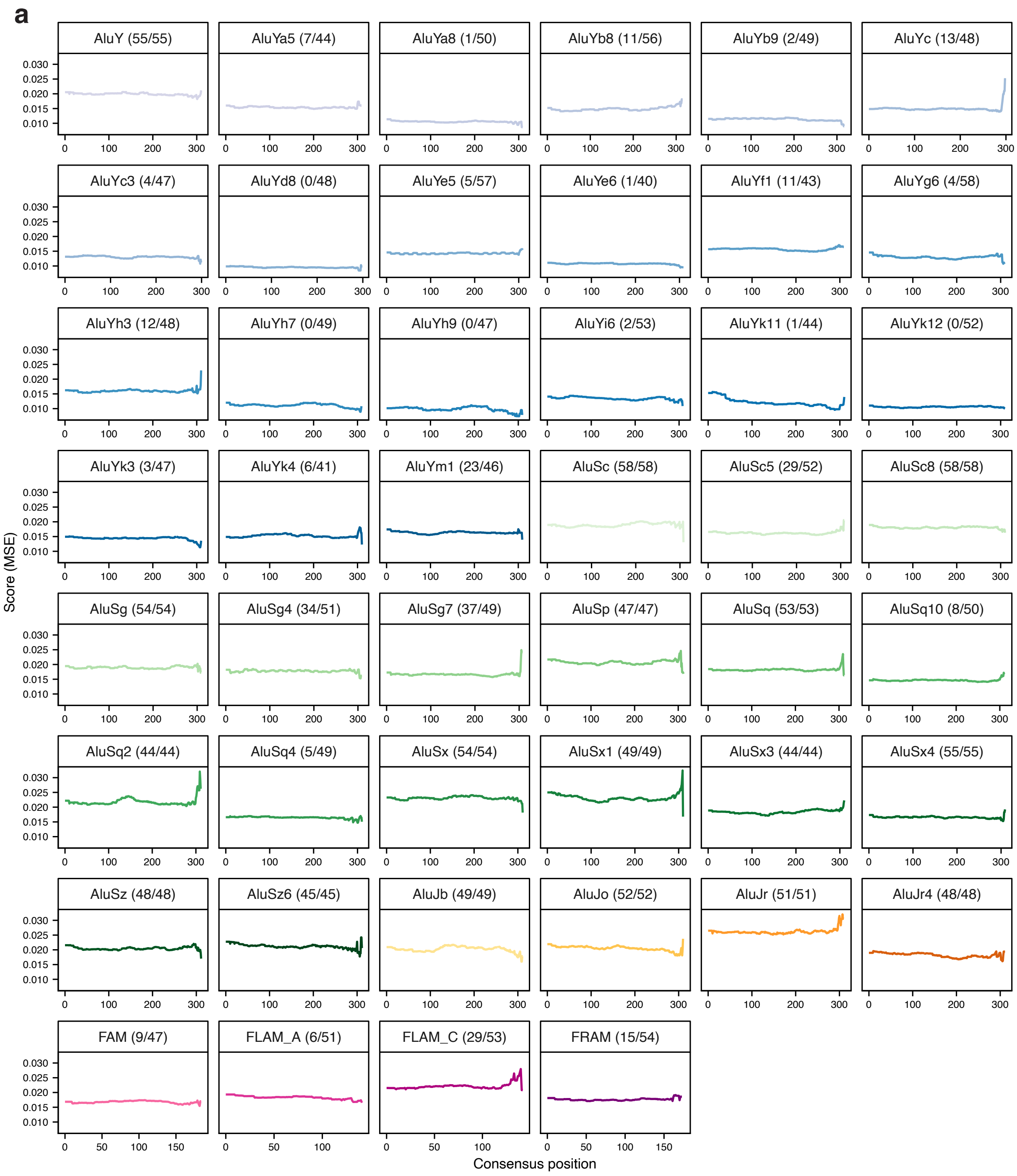


**Supplementary Figure 13. Tiled deletions across *Alu* elements. a.** Average predicted disruption following deletion of 30-bp windows tiled (stride=10bp) across *Alu* elements, mapped to the corresponding positions in the *Alu* consensus sequence. Values represent the average effect at each position across the 100 most disruptive *Alus* from each of 47 *Alu* families in the dfam database. Only *Alus* on the positive strand are shown. Numbers indicate the number of highly disruptive *Alus* represented for each family.

### Supplementary Figure 14

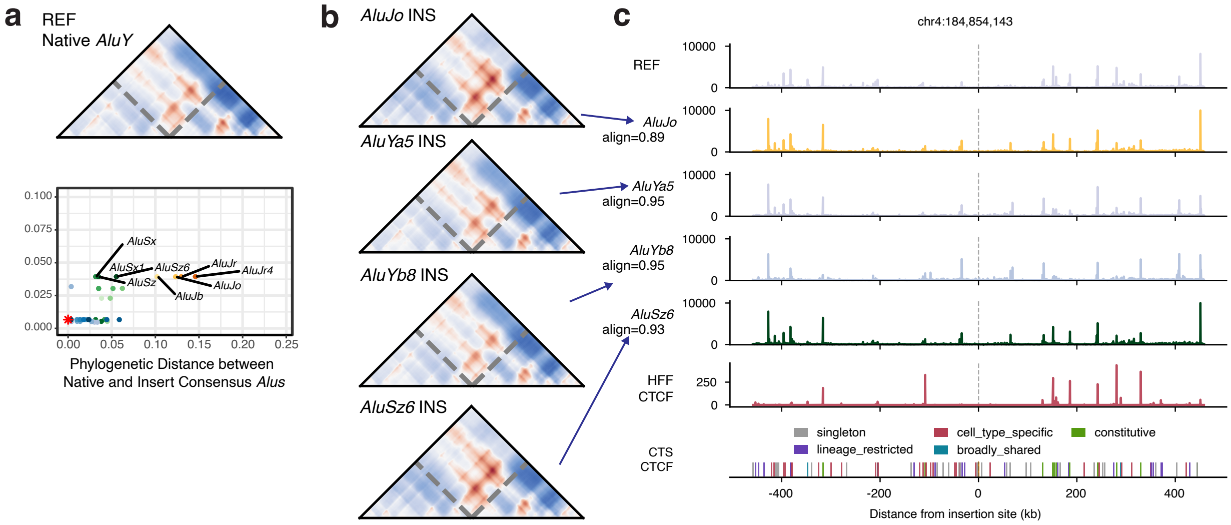


**Supplementary Figure 14. Insertions of consensus *Alu* sequences at an *AluY* locus on chr4. a.** Predicted contact map for a native *AluY* at chr4:184,853,990–184,854,296 (top) and MSE between the predicted contact maps of the native *AluY* and each inserted consensus *Alu* versus phylogenetic distance from the native *AluY* (bottom). **b.** Predicted contact maps following insertion of consensus Alu sequences from different subfamilies at the locus shown in (**a**). **c.** Input saliency for sequences containing the inserted Alu sequences shown in (**b**).

### Supplementary Figure 15

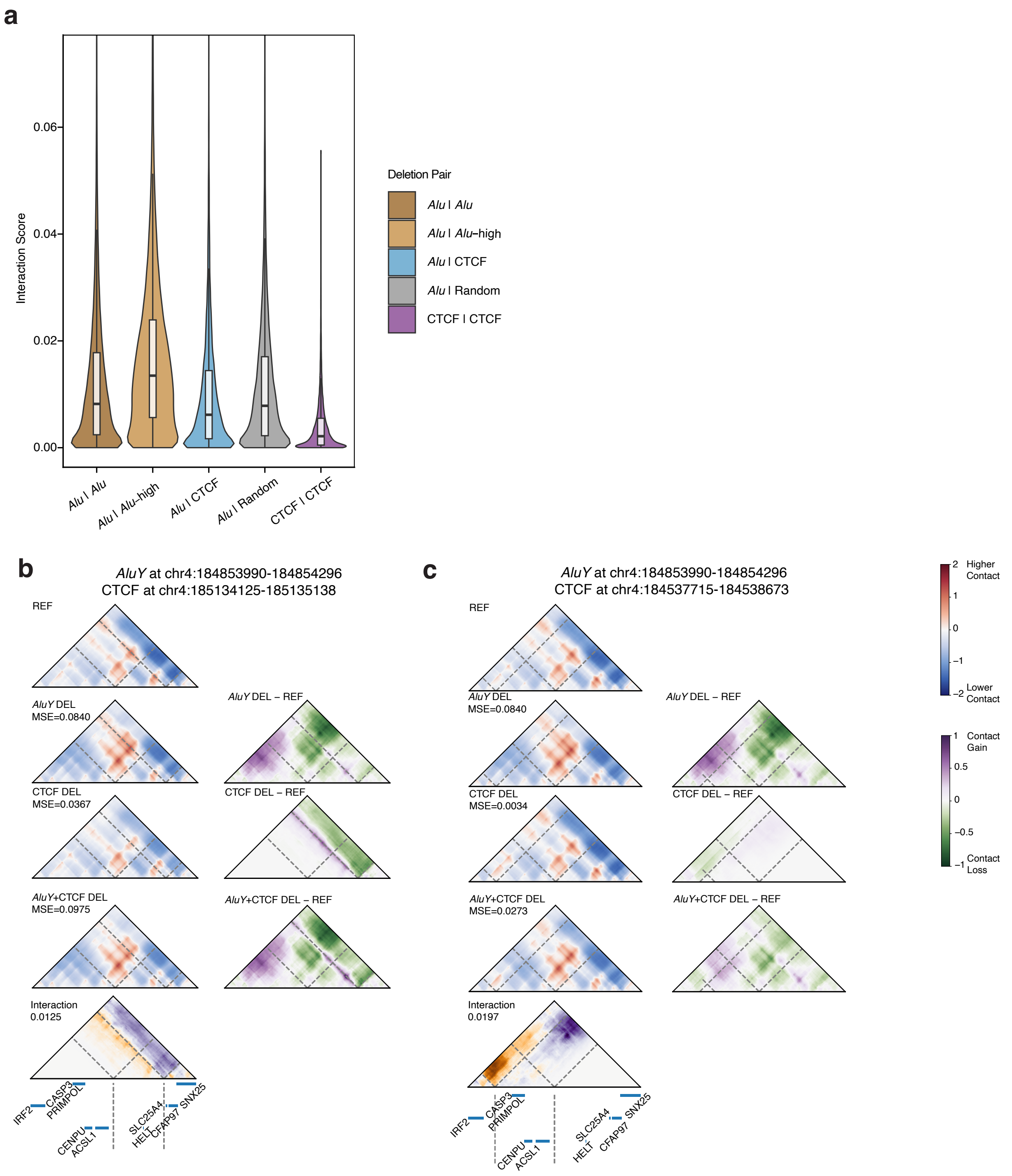


**Supplementary Figure 15. Non-additive interactions between *Alus* and nearby CTCF sites. a.** Distribution of interaction scores for paired deletions of *Alu*–*Alu*, *Alu*–disruptive *Alu*, *Alu*–CTCF, *Alu*–random, and CTCF-CTCF pairs across 1,000 randomly sampled highly disruptive *Alus*. **b-c.** Predicted contact maps following deletion of two distinct CTCF sites near an *AluY* element on chr4. Predicted contact maps follow deletion of the *AluY* element, each CTCF site individually, and the *AluY* and CTCF sites in combination.

### Supplementary Tables

**Supplementary Table 1.** Predicted SuPreMo-Akita disruption scores for all *Alu* elements in the hg38 reference genome, annotated with genomic and epigenomic features.

**Supplementary Table 2.** Predicted disruption scores for polymorphic SVs from the 1000 Genomes Project long-read call set.

**Supplementary Table 3.** Predicted SuPreMo-Akita disruption scores for all *Alu* elements in the T2T-CHM13 reference genome.

**Supplementary Table 4.** Performance metrics for regression models predicting *Alu* disruption scores from genomic annotations.

**Supplementary Table 5.** Predicted disruption scores from *in silico Alu* insertion swaps performed at matched loci within individual genes.

**Supplementary Table 6.** Predicted disruption scores for 1,000 randomly sampled high-disruption and 1,000 neutral *Alu* elements, for ISM experiments in Figures 5-6. These include random nearby deletions, *Alu* nucleotide shuffling, GC/CpG content mutagenesis, consensus *Alu* insertions, and pairwise element deletions.

**Supplementary Table 7.** Enrichment results for transcription factor binding motif overlap in high- versus low-disruption *Alu* elements.
